# The Dynamics of the Female Microbiome: Unveiling Abrupt Changes of Microbial Domains across Body Sites from Preconception to Perinatal Phase

**DOI:** 10.1101/2023.08.31.555744

**Authors:** Charlotte J Neumann, Manuela-Raluca Pausan, Victoria Haid, Eva-Christine Weiss, Vassiliki Kolovetsiou-Kreiner, Bettina Amtmann, Petra Winkler, Alexander Mahnert, Evelyn Jantscher-Krenn, Christine Moissl-Eichinger

## Abstract

The microbial ecosystem of women undergoes enormous changes during pregnancy and the perinatal period. Little is known about the extent of changes in the maternal microbiome beyond the vaginal cavity and its recovery after birth. In this study, we followed pregnant women (mpre, *n* = 30) into the postpartum period (1 month postpartum, mpost, *n* = 30). We profiled their oral, urinary, and vaginal microbiome, archaeome, mycobiome and urinary metabolome and compared them with nonpregnant women (np, *n* = 29).

Overall, pregnancy status (np, mpre, mpost) had a smaller effect on the microbiomes than body site, but massive transitions were observed for the oral and urogenital (vaginal and urinary) microbiomes. While the oral microbiome fluctuates during pregnancy but stabilizes rapidly within the first month postpartum, the urogenital microbiome is characterized by a major remodeling caused by a massive loss of *Lactobacillus* and thus a shift from vaginal community state type (CST) I (40% of women) to CST IV (85% of women). The urinary metabolome rapidly reached an np-like composition after delivery, apart from lactose and oxaloacetic acid, which were elevated during active lactation. Fungal and archaeal profiles were indicative of pregnancy status. *Methanobacterium* signatures were found exclusively in np women, and *Methanobrevibacter* showed opposite behavior in oral cavity (increased) and vagina (decreased) during pregnancy.

Our findings suggest that the massive remodeling of the maternal microbiome and metabolome needs more attention and that potential interventions could be envisioned to optimize recovery and avoid long-term effects on maternal health and subsequent pregnancies.

## Introduction

The female body undergoes profound changes from conception through pregnancy and the perinatal period that affect hormonal status, metabolism, the immune system, and the microbiome ^1^. For example, the maternal microbiome changes substantially during pregnancy, a period characterized by low-grade inflammation, increased fat storage, and insulin resistance ^2^. However, knowledge of the transition of the microbiome from nonpregnant to pregnant and especially from prepartum to postpartum is relatively sparse, particularly with respect to bacteriomes, mycobiomes, and archaeomes, and beyond vaginal and gastrointestinal body sites.

It is well known that the vaginal microbiome is significantly altered by pregnancy. In general, a healthy vaginal microbiome is dominated by lactobacilli, which are thought to maintain low pH and produce bacteriocins that inhibit pathogen growth^3^. During pregnancy, vaginal diversity decreases and the dominance of *Lactobacillus*-species increases ^4^, probably to protect mother and fetus from infection ^5^. This rise in *Lactobacillus* may be due to increased estrogen levels during pregnancy, which cause maturation of the vaginal epithelium and consequent accumulation of glycogen. Degraded by the host, products such as maltose, maltotriose and maltotetraose then promote the growth of lactic acid bacteria ^6,7^. Pregnancy also affects the oral and urinary microbiome. During pregnancy, the oral microbiome is of particular clinical interest, as many women suffer from bleeding gums, gingivitis or periodontitis ^8,9^, which are likely caused by microbial disturbances associated with immune modulation and hormonal changes ^10^. Gingival diseases are associated with adverse pregnancy outcomes ^11^. Similarly, the urinary microbiome of pregnant women is clinically relevant because urinary tract infections increase the risk of preterm birth ^12^. A previous study showed that certain microbial taxa in urine, namely *Ureaplasma urealyticum,* were associated with preterm birth even in the absence of urinary tract infections ^13^.

Parturition marks a rather abrupt change for both child and mother, which also brings dramatic changes in microbial niches. For the newborn, birth is a starting point for microbial colonization. The transfer of the microbiome from mother to child plays an essential role in the development of the infant’s microbiome and immune system. For the mother, childbirth is a turning point that also affects her microbiome transitioning from a pregnant to a postpartum state. In addition, the immediate postpartum period is a difficult time, characterized, for example, by drastic hormonal changes, increased energy and nutrient demands (also due to breastfeeding), sleep disturbances, or depressive symptoms ^14,15^, which can affect the microbiome and *vice versa* ^15^.

After delivery, the vaginal microbial community shifts from a *Lactobacillus-*dominated microbiome to a *Lactobacillus*-depleted microbiome ^15^, likely caused by the rapid decline in estrogen levels^3^. This phase is associated with diseases such as vaginal bacteriosis and vulvovaginal candidiasis ^16,17^. Cultivation-based studies have shown that *Gardnerella, Peptococcus, Bacteroides, Staphylococcus, Streptococcus* and *Ureaplasma* species ^3^ are increased associated with postpartum endometritis, which occurs in 1-3% of all deliveries ^18^. It is estimated that only about 50% of women achieve a healthy *Lactobacillus*-dominated status at one year postpartum because *L. crispatus* does not return to predominance ^14^.

Maternal gut microbiota has been shown to be altered postpartum ^19^. In addition, pregnancy complications such as gestational diabetes can cause perturbations in the composition of the gut microbiome during pregnancy that can have lasting effects after delivery ^15^.

Although awareness of the problematic recovery of the vaginal microbiome to the pre-pregnancy status quo has been raised, potentially impacting subsequent pregnancies ^14^, knowledge of the transition of the maternal microbiome of also other body sites as well as the dynamics of the non-bacterial microbiome is still surprisingly sparse as already indicated elsewhere^19^. Here we highlight the transition of microbiomes in the oral and urogenital (urinal and vaginal) body sites of 30 women from the prepartum to one month postpartum and place it in context with the microbiomes of nonpregnant women. We use specific detection and analysis methods for the archaeome and mycobiome to provide a broader view of the holistic microbiome.

We found that archaea and fungal representatives are also affected by the massive remodeling of the microbiome during pregnancy and that archaeal signals from the genus *Methanobrevibacter* reflect vaginal niche restriction and oral niche opening during pregnancy. Overall, unlike the vaginal microbiome, the oral microbiome recovers rapidly after delivery and reaches pre-pregnancy status, whereas vaginal dysbiosis after delivery is long-term and accompanied by a massive shift in the urinary microbiome and metabolome.

## Results

### Study set-up and general description

The aim of the study was to understand the extent of imbalance of the bacterial and non-bacterial, oral, gut and urogenital (urinal and vaginal) microbiomes in postpartum women compared to their pregnant status or nonpregnant individuals. Thirty healthy pregnant participants were enrolled in this study to investigate the microbial changes during and after pregnancy. For this purpose, we recruited women in the third trimester (*n* = 30, “mpre”), who provided samples at this time and one month after delivery (*n* = 30, “mpost”; Cesarean section (CS) *n* = 15, vaginal delivery *n* = 15). The study also included nonpregnant female controls for comparison (*n* = 29, “np”). The following samples were collected: Stool (np, mpre), oral swabs (all), non-cathetered urine (all), vaginal swabs (all). A total of 353 samples were processed by amplicon sequencing targeting the overall microbiome (“universal”), archaea and fungi. NMR-based metabolomics was performed for the urine samples. An overview on the samples, number of reads, and Amplicon Sequence Variant (ASV) numbers are shown in Supplementary Fig. 1 and 2. The characteristics of the cohorts are shown in Table 1.

**Table 1:** Characteristics of the study cohort, statistically compared between the pregnant group pre- and postpartum (mpre and mpost) and the nonpregnant (np) group as well as within the pregnant group between those who delivered via Cesarean section (CS) versus those who delivered vaginally.

| Characteristics | mpre & mpost<br>(n = 30) | np<br>(n = 29) | P-value<br>(1) T-test<br>(2) Mann Whitney U test<br>(independent samples)<br>(3) Fisher's exact t-test |
| --- | --- | --- | --- |
| Mean age (yr) ± SD (range) | 34.5 ± 4.7 (27 - 44) | 32.0 ± 5.7 (19 - 43) | 0.083<sup>(1)</sup> |
| Mean BMI prepregnancy (kg/m2)<br>± SD (range) | 22.4 ± 4.1 (17.3 - 37.0) | 22.6 ± 3.6 (17.2 - 31.5) | 0.968<sup>(2)</sup> |
| Pregnant group |  |  |  |
|  | CS (n = 15) | Vaginal birth (n = 15) | p-value |
| Mean age (yr) ± SD (range) | 35.8 ± 5.1 (28 - 44) | 33.1 ± 3.9 (27-39) | 0.113<sup>(1)</sup> |
| Mean BMI at sampling (kg/m2) ±<br>SD (range) | 27.3 ± 6.2 (21.8 - 47.6) | 27.6 ± 3.4 (22.2 - 34.0) | 0.436<sup>(2)</sup> |
| Mean BMI prepregnancy (kg/m2)<br>± SD (range) | 22.2 ± 4.5 (18.3 - 37.0) | 23.0 ± 3.8 (17.3 - 29.4) | 0.285<sup>(2)</sup> |
| SS-days | 272.7 ± 7.3 (258 -286) | 283.1 ± 3.5 (276 -289) | < 0.001<sup>(1)</sup> |
| Infant weight (g) | 3408.6 ± 465.5 (2510 - 4250) | 3486.0 ± 269.8 (3040 - 4110) | 0.582<sup>(1)</sup> |
| Days in hospital | 3.9 ± 0.7 (2 -5) | 3.6 ± 1.5 (1 - 6) | 0.87<sup>(2)</sup> |
| Sex of the infant | F: 8; M: 7 | F: 6; M: 9 | 0.715<sup>(3)</sup> |

### Bacterial and fungal microbiomes are affected by pregnancy status

Differentiation of the bacterial microbiomes was mainly determined by body site (Fig. 1A; permanova, method = “bray”, *p* = 0.001, *R*^2^ = 0.194), and, with a smaller effect size, also by maternal pregnancy status (mpre, mpost, and np; permanova, method = “bray”, *p* = 0.001, *R*^2^ = 0.03). In all samples, *Lactobacillus* was the most abundant genus, found primarily in vaginal and urinary np and mpre samples, followed by *Streptococcus* (mostly oral samples), and *Gardnerella* (urogenital samples) (Fig. 1B).

**Figure 1:**
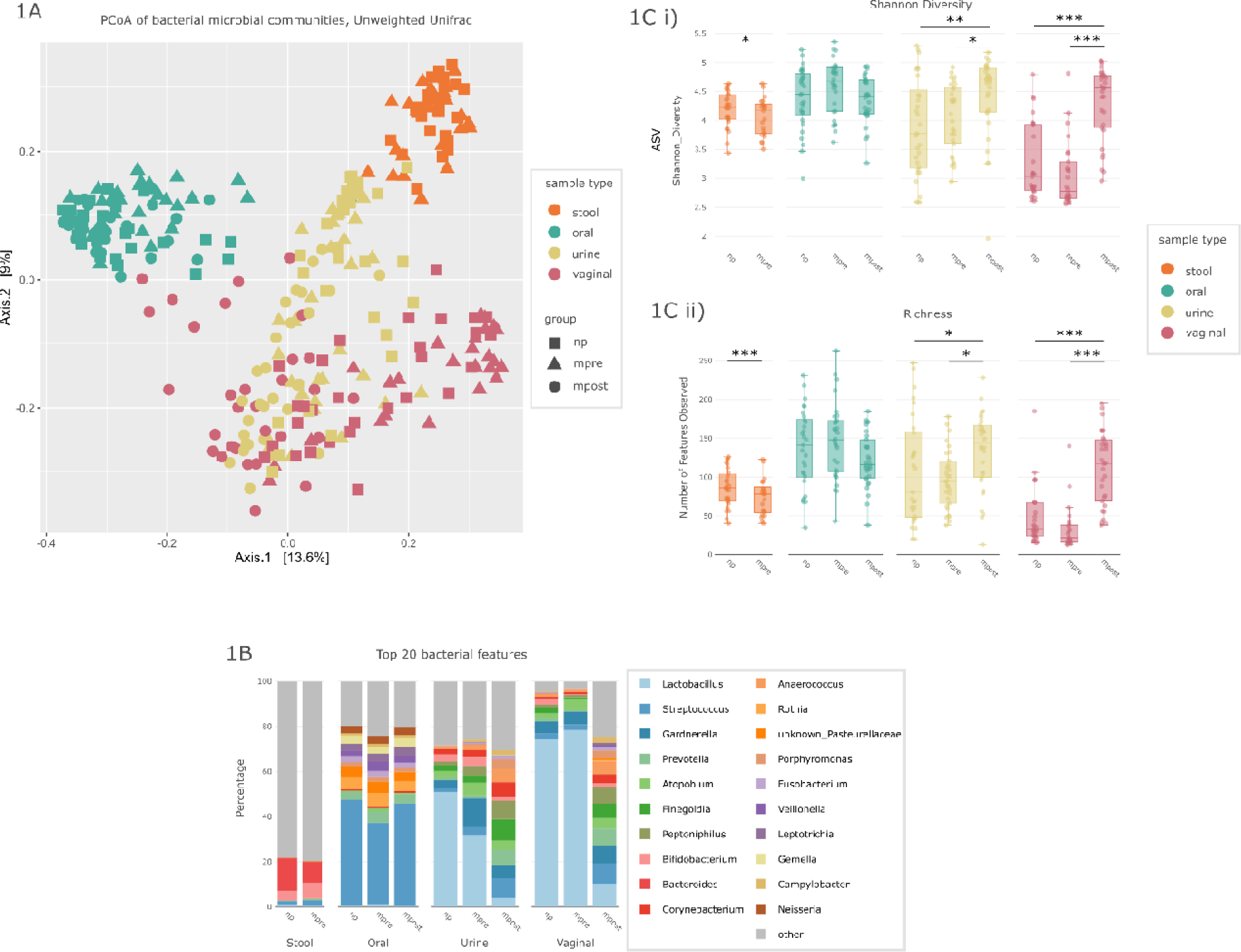
Beta diversity, alpha diversity and composition of the bacterial microbiome. A) Principle Coordinate Analysis (PCoA) of sample type (color) and group (shape) with Unweighted Unifrac distance matrix on Amplicon Sequence Variant (ASV) level. B) Alpha diversity of sample types by group of i) Shannon diversity and ii) richness. C) Composition on the top 20 bacterial genera in relative abundance, depicted by sample type and group. Significance levels are indicated with asterisks for *p*l<l0.001 (**), *p*l<l0.01 (**), *p*l<l0.05 (*).

The impact of pregnancy (mpre compared to np controls) on the fecal bacteriome was significant for alpha diversity (Fig. 1C, ASV level) but rather low on taxon level (Suppl. Fig. 3). This was reflected by the fact that no statistically significantly different abundances of bacterial genera, species or ASVs were found after Benjamini Hochberg (BH) correction (Suppl. Fig. 4). Overall, pregnant women had fewer *Bacteroides* signatures but relatively more *Bifidobacterium* in their gut microbiome, as reported in other studies ^20^ (Fig. 1B). The gastrointestinal archaeome was not affected by pregnancy.

However, pregnant women carried less fungal signatures in their gastrointestinal tract than nonpregnant women, as success in amplifying ITS regions in mpre samples was lower (n = 5/30, compared to n = 20/29 in np and n = 28/30 in mpost) (Suppl. Fig. 2). Nevertheless, the composition of the fecal mycobiome was quite uniform between the groups for which data were available (Fig. 2A). *Candida* and *Saccharomyces* were the most abundant fungal genera (Suppl. Fig. 6), similar to previous findings ^21^. The only significantly differentially abundant signatures were unclassified fungal signatures at species level before BH correction.

**Figure 2:**
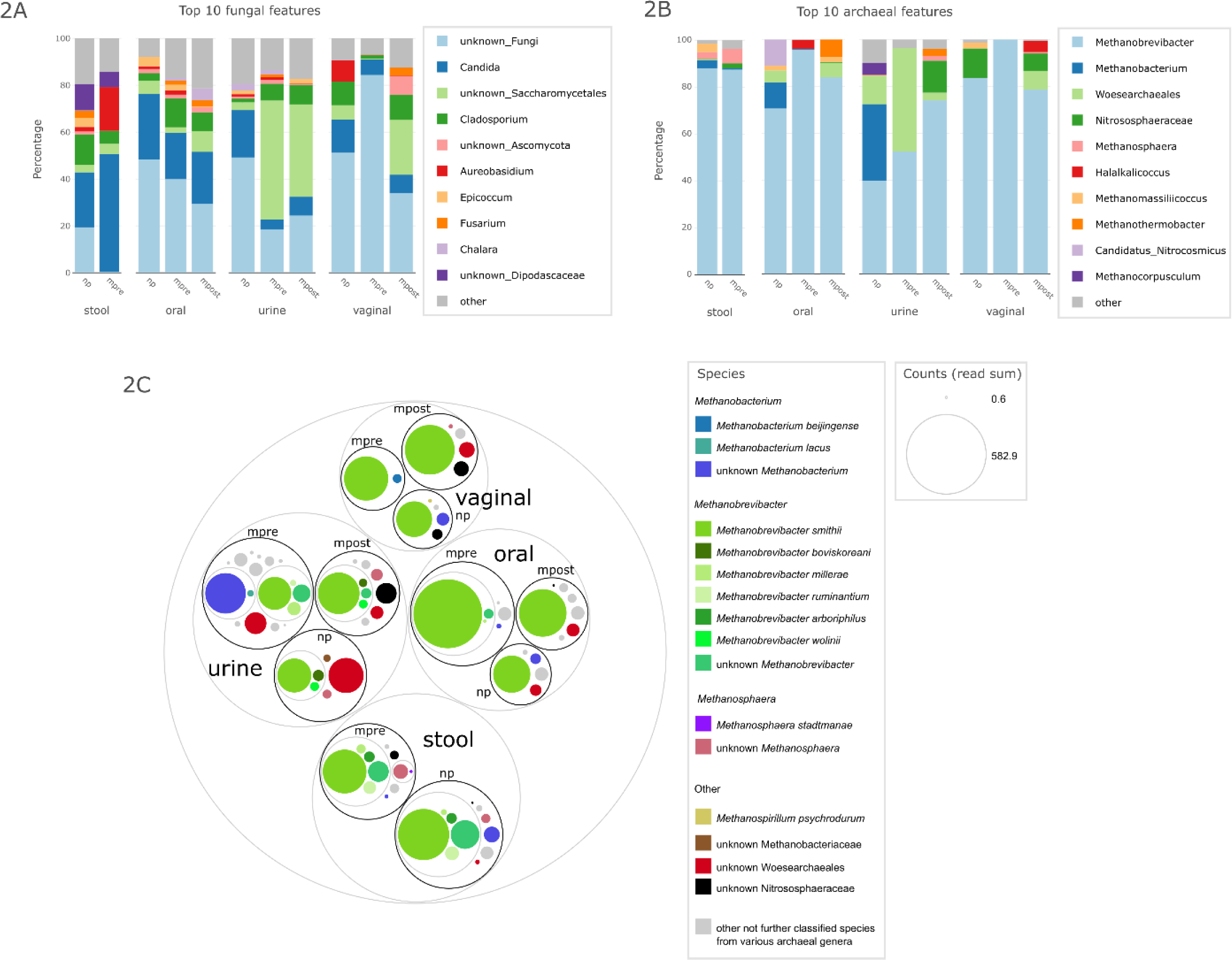
Relative abundance of top 10 A) fungal genera and B) archaeal genera, depicted by relative abundance, depicted by sample type and group. C) circle packing plot of archaeal species depicted by body site and group; the size of the colored dots represents counts of normalized read sums.

### Methanogenic archaea - indicators for pregnancy?

Archaeal signatures were analyzed semi-quantitatively (“universal approach”) alongside with bacteria and qualitatively in a separate “nested approach” specifically targeting archaea.

Overall, archaeal richness was very low in all samples from all body sites (max. four genera/ six species/ 29 ASVs per individual and body site, Suppl. Fig. 7). A total of 28 archaeal genera, 38 species and 411 ASVs were detected in the data set. *Methanobrevibacter* was the most prevalent archaeal genus in samples from all body sites (Fig. 2B), followed by *Methanobacterium*, which not found in mpost group but in all sample types, especially in urine but also in oral, vaginal and stool samples (Fig. 2C).

The most abundant *Methanobrevibacter* species was *M. smithii* which represented the three most abundant ASVs (ASVs 1, 8, 9). These three ASVs accounted for 58.25% of all archaeal reads, 77.18% of *Methanobrevibacter* reads and 88.01% of all *M. smithii* reads (Fig. 2C). Known as a gut-associated archaeon, *M. smithii* indeed dominated the gastrointestinal archaeome (67.83% of all archaeal reads in the np and mpre groups). Moreover, the fecal samples showed the highest richness of *Methanobrevibacter* species and ASVs than the other sample types (Fig. 2C). It should be noted that due to insufficient databases for the classification of human-associated archaea, species classification is likely not comprehensive and therefore species information should be taken with caution ^22^. Classification of *Methanobrevibacter* in particular ws further complicated by the fact that the two most common species, *M. smithii* and *M. intestini*, cannot be distinguished using 16S rRNA gene amplicons ^23^.

*Methanobrevibacter* species were also found in urine samples and in the vagina and oral cavity.

The urinary archaeome did not significantly differ between the groups (Unweighted Unifrac, Adonis, *p* = 0.001). We observed a decrease in Shannon diversity and richness of archaea from np to mpre and mpost (Suppl. Fig. 7).

The vaginal archaeome carried a variety of archaeal signatures, but overall had the lowest Shannon diversity and richness of archaeal genera, species and ASVs. Mpre samples contained exclusively *Methanobrevibacter (smithii,* ASVs 1,8,9), in contrast to np and mpost samples, which contained other archaeal signatures such as unclassified Nitrososphaeraceae and Woesearchaeales, *Halalkalicoccus*, *Methanomassiliicoccus*, *Methanosphaera* and *Methanospirillum*) (Suppl. Fig. 8). Interestingly, only few mpre women exhibited any archaea in their vaginal microbiome, and when they did, they were exclusively *Methanobrevibacter (smithii)* (Suppl. Fig. 9). Although the presence of archaea in the vaginal microbiome increased after delivery, no statistically significant differences in the Principal Coordinate Analysis (PCoA) were observed between the groups (Suppl. Fig. 5).

In contrast, the oral samples revealed a significant increase in the number (richness) and diversity (Shannon) of archaeal ASVs in mpre compared to np and mpost. This was observed not only in the archaea specific amplicon approach indicating relative abundance, but also in the universal approach for which differential abundance could be calculated. *M. smithii* was found significantly increased in mpre compared to mpost (Aldex2, *p* = 0.005, *q* = 0.321) and to np (Aldex2, *p* = 0.003, *q* = 0.300). It should be mentioned that, surprisingly, the obtained sequences indeed were classified as *M. smithii* and not as *M. oralis* although classifications should be taken with caution due to incomplete databases (see above).

In fact, not a single np woman carried *M. smithii* in her oral cavity, but 16 prepartum women did. After delivery, *M. smithii* was detected in only one mpost woman (Suppl. Fig. 11). We therefore suggest that the oral niche of a pregnant woman favors the growth of the anaerobic methane-producing archaea *Methanobrevibacter*, and *M. smithii* in particular.

In summary, we observed niche restriction for methanogenic archaea in the vagina during pregnancy, whereas the niche in the oral cavity was opened. This indicates that *Methanobrevibacter* species may reflect changes in the ecosystem.

### The oral bacteriome and mycobiome undergoes temporary changes during pregnancy

In the oral samples, alpha diversity of the bacteriome did not change substantially with pregnancy status (Fig. 1C). However, beta diversity significantly differed between the groups, as shown by PCoA plots (Unweighted Unifrac, *p* = 0.001 ANOVA; Fig. 1A, Suppl. Fig. 10). It is noteworthy that the mpre samples clustered visibly apart from the other sample groups. We hypothesize that the differences between the groups are mainly due to rarely abundant ASVs, such as the numerous, highly divergent *Streptococcus* ASVs. *Streptococcus* was identified as the most abundant genus in all three groups, accounting for 43.08% of relative abundance (Suppl. Fig. 3).

By performing differential abundance analysis, we did not identify a single genus, species or ASV (with the exception for archaeal *Methanobrevibacter* (*smithii*), see above) that was differentially abundant between mpost and np (with or without BH correction; Suppl. Fig. 4), indicating that a similar microbial status to pre-pregnancy was achieved relatively quickly after pregnancy (see Fig. 1C). However, the mpre samples showed a variety of differences (Suppl. Fig. 4). However, none of these differences remained significant after BH correction, so only trends can be reported (Aldex2, *p* < 0.05, *q* > 0.05). As mentioned above, *Methanobrevibacter* signatures, particularly those of *M. smithii,* were specifically increased in mpre oral samples compared to mpost (*p* = 0.0047, *p* = 0.321) or np (*p* = 0.003, *q* = 0.300). This trend was opposite to that observed for *Streptococcus*. Overall, 13 streptococcal ASVs were increased in the np group and six in the mpost group compared with the mpre samples (*p* < 0.050, Suppl. Fig. 4). Furthermore, the relative abundance of *Streptococccus* ASVs in mpre was rather individual and less uniform than in np and mpost, as shown by the wider range of the confidence interval (Suppl. Fig. 4).

It has been reported that the incidence of periodontal disease increases during pregnancy ^24^, possibly leading to adverse pregnancy outcomes such as preterm birth ^25^. In our mpre group, we did not observe a significant increase in the so-called orange and red complex bacteria, which have been shown to accountfor initiation and progression of periodontal disease ^26^ (Suppl. Fig. 12 and Suppl. Fig. 13). It shall be mentioned though, that metadata on participants’ dental status was not available, not allowing further interpretation. However, *Methanobrevibacter* load or presence has been previously associated with severe periodontitis ^27,28^.

Fungal diversity in the oral cavity increased from np to mpre (Suppl. Fig. 14). Highly statistically significant differences (Suppl. Fig. 15, Unweighted Unifrac, Adonis, *p* = 0.001) between the three groups were observed by Adonis. These strong differences can be explained solely by the highly significant differences in two fungal ASVs (ASVs 8 and 9) that were not further classified. ASV8 was highly significantly increased in np compared to mpre (Aldex2, *p* < 0.001, *q* < 0.001) and to mpost (Aldex2, *p* < 0.001, *q* < 0.001), whereas ASV9 was significantly more abundant in mpre than in np (Aldex2, *p* = 0.030, *q* = 0.394). Overall, approximately 50% of all oral fungal samples could not be classified. The main members of the oral mycobiome were *Candida albicans* and *Cladosporium cladosporioides* followed by not further classified Saccharomycetales, which were shared between the three groups.

Overall, we observed temporary fluctuation of the oral microbiome during pregnancy and rapid stabilization of the oral bacteriome and archaeome thereafter.

### The urogenital mycobiome and bacteriome undergo vast changes during and after pregnancy

Due to the local proximity of the urinary and vaginal tracts, an overlap can be biologically expected. We specifically collected midstream urine to largely prevent transfer during sampling. Nevertheless, the overlap was substantial: 77 genera in np, 47 in mpre and 125 in mpost were shared (Table 2).

**Table 2:** shared and unique bacterial genera between vaginal and urine plotted per group.

|  | Urine unique | Shared urine and vaginal | Vaginal unique | total |
| --- | --- | --- | --- | --- |
| np | 470 | 77 | 18 | 565 |
| mpre | 302 | 47 | 9 | 358 |
| mpost | 174 | 125 | 56 | 355 |

To numerically assess the microbial exchange between the two sites in the context of all other body sites sampled, we performed source tracking analysis (Fig. 3A, Suppl. Fig.16). In all samples, the contribution of the oral microbiome towards the urogenital tract was marginal and did not increase during pregnancy. Overall, the exchange of vaginal and urinary microbial patterns was very clear (Fig. 3A). Notably, the urinary microbiome in mpost contributed significantly less to the vaginal microbiome than in mpre samples (Wilcoxon Rank, mpre: 95.25% (median), mpost: 37.18 % (median), *p* = 0.002 Suppl. Fig. 16). Similarly, we observed a trend towards an overall lower contribution of vaginal samples to urine in samples from mpost (t-test, np-mpost, *p* = 0.078, median np: 55.38%, mpre: 38.80%, mpost: 41.54%).

**Figure 3:**
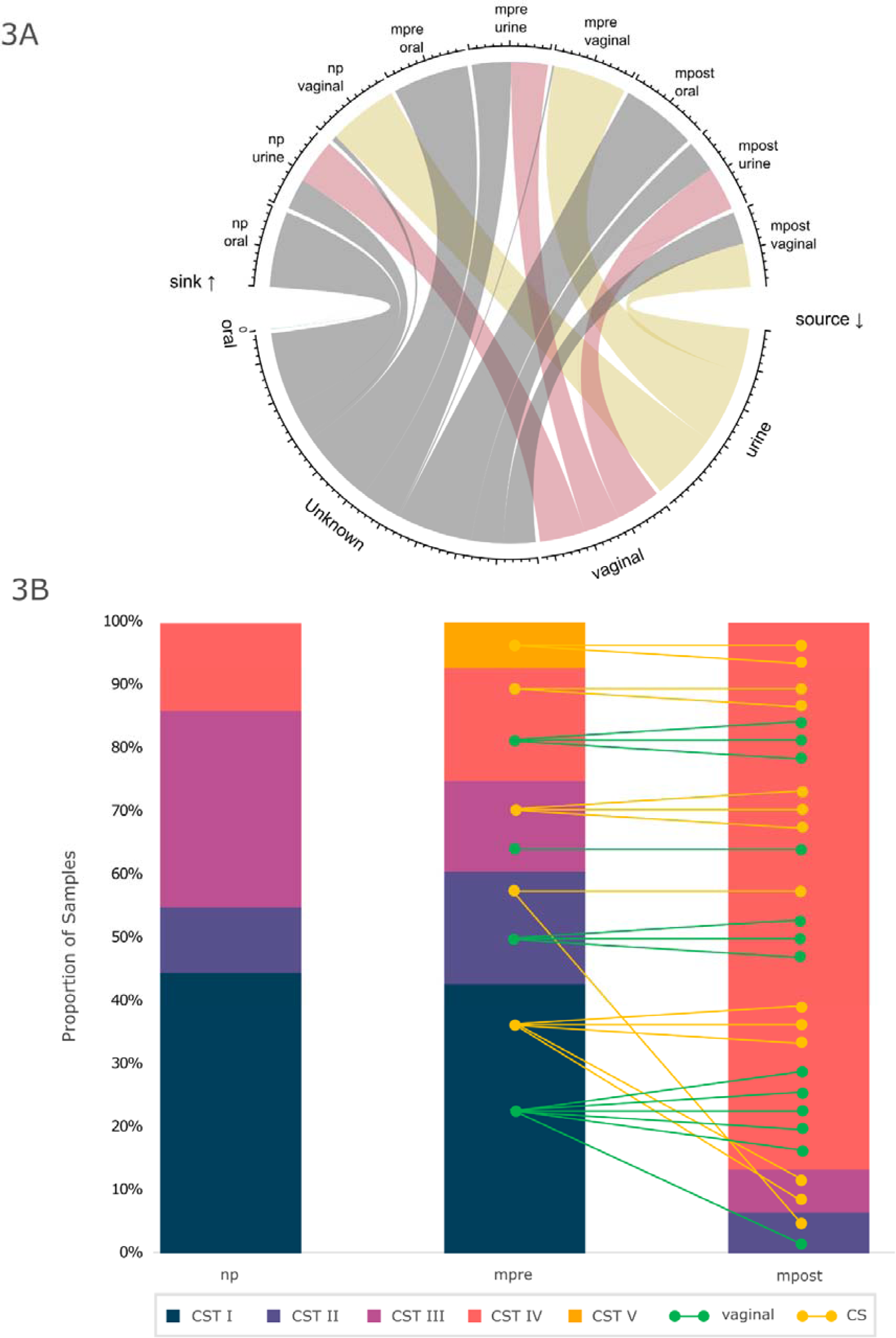
A) Source tracking; comparison of oral, vaginal and urine samples with respect to their contribution from oral, urine, vaginal and unknown sources. B) Vaginal community state types (CST), split per group, lines indicate individuals, colors indicate delivery type; yellow: Cesarean section (CS), green: vaginal

Furthermore, vaginal and urine (urogenital) samples were analyzed comparatively with respect to their mycobiome and overall microbiome.

For both the vaginal and the urinary mycobiome, the groups clustered statistically significantly (Unweighted Unifrac, Anova, *p* = 0.001) on ASV level in the PCoA plots (Suppl. Fig. 15, Suppl. Fig. 5). The vaginal mycobiome, especially in mpre samples, was dominated by reads that could not be fully classified (Suppl. Fig. 6, Suppl. Fig. 17). This was accompanied by a significant decrease in alpha diversity and richness in mpre (mpre-mpost ; Shannon *p* = 0.010, richness *p* = 0.035) (Suppl. Fig. 14) and led to a significant clustering of the groups (Anova, *p* < 0.001, Suppl. Fig. 15). Fungal composition was quite different between groups, but only in relative abundance (Suppl. Fig. 6) and not in differential abundance testing with Aldex2 (Suppl. Fig. 17). Similar to the vaginal mycobiome, the urinary mycobiome was dominated mainly by not further classified fungal patterns, but specifically not further classified Saccharomycetales and Candida (Suppl. Fig. 6) as already described before ^29^. Not further classified ASVs were significantly more abundant in np than in mpre and mpost (Aldex2, np-mpre *q* = 0.044, np-mpost *q* = 0.161) (Suppl. Fig. 17). Diversity, and especially richness decreased sharply from np to mpre (Shannon *p* = 0.035, richness *p* < 0.001) (Suppl. Fig. 14).

Significant changes in alpha diversity were also observed in bacteria (Fig. 1C, ASV level). Urogenital samples were characterized by a significant increase in Shannon diversity from mpre to mpost. Urine samples showed a continuous increase in alpha diversity from np to mpre to mpost. We observed that in some urine samples, bacterial richness was remarkably high in the np group, with some individuals carrying up to ∼150 different bacterial genera. In contrast, vaginal samples had low overall richness (max. 55 different bacterial genera) and diversity. The lowest diversity was observed in vaginal mpre samples, before reaching a higher alpha diversity after birth than in the nonpregnant state (Fig. 1C).

### Delivery has a substantial impact on the urogenital bacteriome driven by a massive loss of *Lactobacillus*

As indicated above, in particular the urogenital microbiome of postpartum women differed significantly from that of pregnant and nonpregnant women, in terms of alpha and beta diversity as well as microbiome composition (Fig. 1A,C). Therefore, beyond composition and diversity, we performed more detailed analyses on these urogenital microbiomes including analysis of Vaginal Community State Types (CSTs) and metabolomics.

The vaginal microbiome in postpartum women was very different to that of np or mpre women (Fig. 1A, Unweighted Unifrac; Anova, *p* = 0.001). Postpartum, alpha diversity strongly increased and was characterized by a massive loss of *Lactobacillus* (*q* < 0.001, Suppl. Fig. 4, Fig. 1C). This affected not only the three main *Lactobacillus* species representatives (*L. crispatus, L. iners, L. gasseri*) but also all other detected species (*L. jensenii*, *L. delbrueckii* subsp. *delbrueckii*, *L. amylovorus*, *L. coleohominis*, *L. mucosae*, *L. ruminis* and *L. sanfranciscensis)*. This loss of *Lactobacillus* in pthe postpartum vaginal microbiome has also been described elsewhere ^3^ and allowed colonization of the vaginal body site with new (potentially unnatural and pathogenic) bacteria. Accordingly, postpartum vaginal samples had increased abundance of anaerobic bacteria taxa such as *Prevotella*, *Atopobium*, *Streptococcus*, *Anaerococcus*, *Finegoldia*, *Peptoniphilus* and others, most of which are also described as prominent members of CST IV ^30^ or bacterial vaginosis (BV) ^31^, see Table 3. Women with a CST IV vaginal microbiome may be at higher risk for BV or other vaginal infections ^32,33^.

**Table 3:** Increase or decrease of specific bacterial taxa in the vaginal microbiome in community state type (CST) IV (from ^3^), bacterial vaginosis (BV) (from ^31^) and the dataset. *P*-values and *q*-values are given for Aldex2 comparing nonpregnant (np) with postpartum (mpost) samples.

| organism | CST IV | BV | Dataset, np to mpost |
| --- | --- | --- | --- |
| *Lactobacillus* | decrease | decrease | decrease, $p < 0.001$ ***, $q < 0.001$ *** |
| *Prevotella* | increase | increase | Increase, $p < 0.001$ ***, $q < 0.001$ *** |
| *Dialister* | increase | | Increase, $p = 0.0011$ **, $q = 0.022$ * |
| *Peptoniphilus* | increase | | Increase, $p < 0.001$ ***, $q = 0.0018$ ** |
| *Atopobium* | increase | increase | Increase, $p = 0.0066$ **, $q = 0.077$ |
| *Finegoldia* | increase | | Increase, $p = 0.0075$ *, $q = 0.087$ |
| *Gardnerella* | | increase | Increase, $p = 0.648$, $q = 0.844$ |
| *Mobiluncus* | | increase | Increase, $p = 0.393$, $q = 0.650$ |
| *Bifidobacterium* | | increase | Increase, $p = 0.348$, $q = 0.609$ |
| *Snaetia* |  | increase | x |
| *Streptococcus* | increase | | Increase, $p = 0.014$, $q = 0.131$ |

To further analyze this interesting prepartum to postpartum shift in the vaginal microbiome, we classified the vaginal microbiomes into CSTs (Fig. 3B). The dominant CST in mpre and np was indeed CST I, dominated by *Lactobacillus crispatus* (45% in np, 43% in mpre). In 70% of the pregnant cohort, the vaginal microbiome shifted from a *Lactobacillus-* dominant CSTs prepartum to CST IV postpartum. Postpartum, CST IV was predominant (23 women, 85%), whereas only five women already had CST IV prepartum. Most women with a prepartum CST other than CST IV switched to CST IV after delivery. This postpartum switch to CST IV has been described previously ^3,14^. In the nonpregnant control group only 14% of the participants had CST IV at the time of sample collection.

Of note, delivery mode (CS versus vaginal delivery) and previous deliveries had no effect on the transition of CSTs from mpre to mpost (Chi-Square test *p* > 0.506). Similarly, the delivery mode had no effect on the overall composition of the urinary microbiome postpartum (Permanova; vaginal: *p* = 0.188, urinary: *p* = 0.214).

### Urinary metabolic profiles mirror the transition from pre- to postpartum

Urine analysis is an important diagnostic tool in clinics, not only during pregnancy, as changes in the chemical composition of urine can indicate health problems. We subjected mpre and mpost urine samples to metabolomics analysis to assess metabolic situations and correlate with microbial features. Metabolic profiles were generated using untargeted and targeted NMR-based metabolomics.

When comparing mpre and mpost samples, a significantly higher lactose content was detected in mpost (*q* < 0.001, t-test after log transformation, paired samples), indicating active lactation. Postpartum women who did not breastfeed (*n* = 2) lacked detectable lactose in their urine. In addition, a significant increase in oxaloacetic acid was observed (*q* = 0.011), which might be explained by an altered metabolic status during breastfeeding ^34^. These two compounds were also significantly higher in postpartum samples compared to nonpregnant controls (lactose: *q* < 0.001; oxaloacetic acid: *q* < 0.001; t-test after log transformation, un-paired samples), but no other compounds were found to be significantly different. However, the urine samples from pregnant women had significantly increased levels of, for example, dimethylamine, alanine, glycine, lactic acid and other compounds, as shown in Fig. 4A.

**Figure 4:**
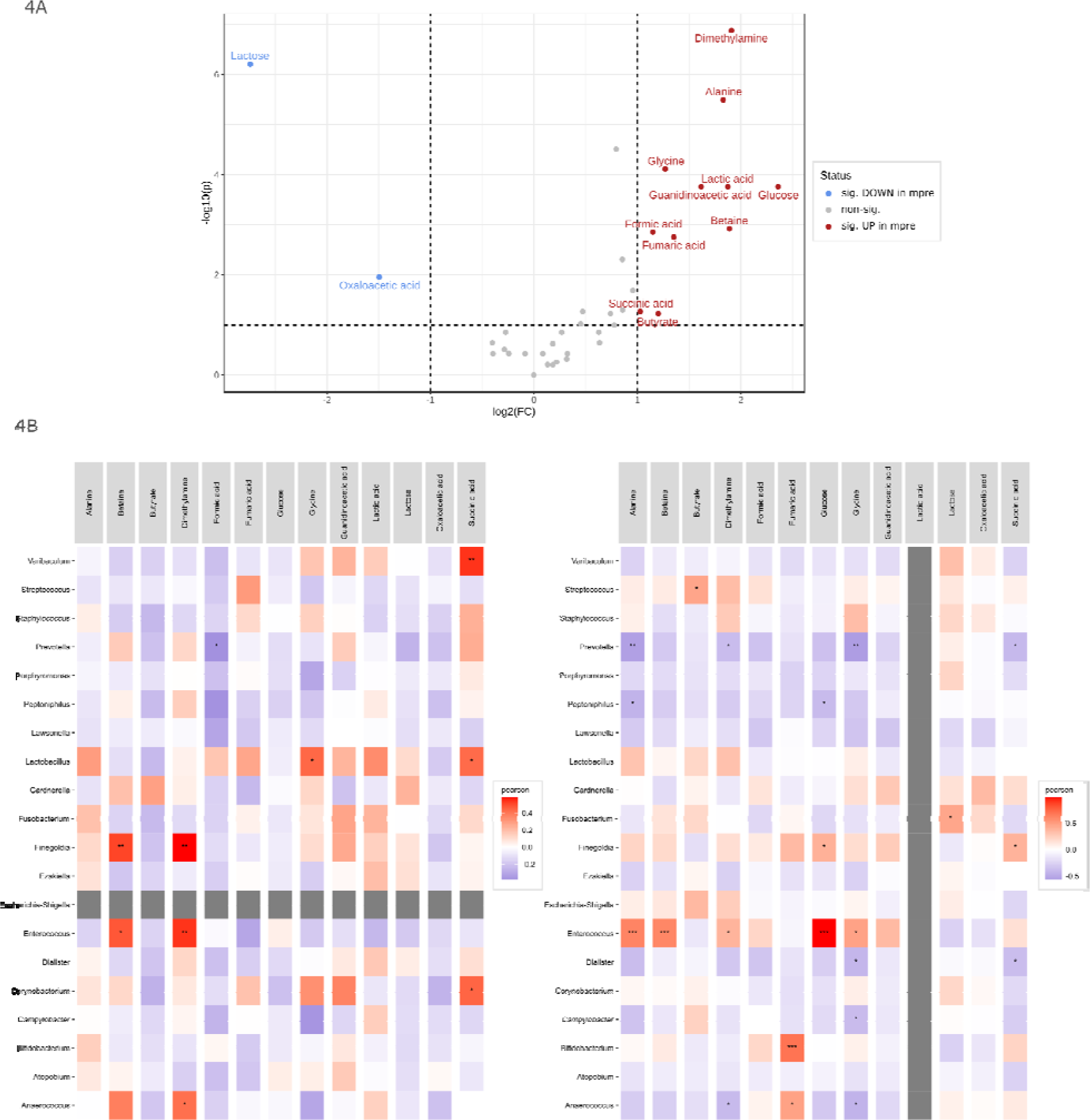
A) Volcano plot of significantly differentially abundant metabolites between mpre and mpost B) Pearson correlation of top 20 abundant bacterial genera with 13 metabolites in urine samples which are differentially abundant between mpre and mpost. left) mpre right) mpost. The grey entries for Escherichia-Shigella in mpre and Lactic acid in mpost reflect absence of those features or metabolites in the subset. Significance levels are indicated with asterisks for pO<O0.001 (***), pO<O0.01 (**), pO<O0.05 (*)

Prepartum, (Fig. 4B i), none of the individuals carried *Escherichia-Shigella*. *Finegoldia*, *Enterococcus* and *Anaerococcus*. Those genera are correlated with betaine and dimethylamine. In addition, we found significant correlations for *Varibaculum* with succinic acid (*q* = 0.003), *Lactobacillus* with glycine and succinic acid, and *Corynebacterium* with succinic acid. Postpartum, (Fig. 4B ii), *Enterococcus* correlated strongly with alanine (*q* < 0.001), betaine (*q* < 0.001) and glucose (*q* < 0.001) as well as with dimethylamine (*q* = 0.047) and glycine (*q* = 0.012). Lactic acid was not detectable in any of the mpost samples from this dataset. Fumaric acid was positively correlated with *Bifidobacterium* (*q* < 0.001) and *Anaerococcus* (*q* = 0.012). *Prevotella*, *Porphyromonas*, *Peptoniphilus* und *Lawsonella* were weakly negatively correlated with all 13 metabolites in both, mpre and mpost.

In the analysis of the whole group (mpre and mpost), the genus *Lactobacillus* strongly correlated with glucose and lactic acid, and was significantly negatively correlated with one of its important substrates, lactose (Suppl. Fig. 18). In healthy situations, the production of lactic acid lowers the pH, which prevents the growth of other microbes and promotes the acid-tolerant lactobacilli. Indeed, lactic acid was reduced in mpost samples (Fig. 4A,Fig. 4B ii), highlighting the loss of acid-producing *Lactobacillus* activity postpartum.

Although the increase of lactose and oxaloacetic acid in urine might be correlated with one of the 20 most abundant microbial genera, lactose in combination with the acid might favor the growth of lactobacilli in the long term.

## Discussion

In this paper, we demonstrated the dynamic transition of the maternal microbiome and urinary metabolome from the preconceptional to the perinatal period. We revealed abrupt changes in the bacterial, fungal, and archaeal components of the microbiome in the oral and urogenital (urine and vaginal) body regions of 30 women from the pregnant state to one month postpartum and placed them in context to the microbiome of nonpregnant women. Understanding the normal postpartum microbiome transition and the influence of maternal (modifiable) factors could help develop prevention strategies for women who tend to develop typical microbiome-associated postpartum health problems or infections.

Pursuing a molecular approach that allowed us to capture nonbacterial components of the microbiome added interesting aspects to the microbiome transition. The overall archaeome profiles found in the urogenital tract (mainly Methanobacteriota and Halobacteriota) were consistent with the literature ^35^ but were not indicative of pregnancy status. Other signals reflected the pregnancy status well: we detected *Methanobacterium* almost exclusively in nonpregnant women (all four body sites examined). Even clearer evidence was provided by the major player among the archaea, *Methanobrevibacter*. The abundance of this archaeon was significantly increased in the oral cavity of pregnant women, but rapidly decreased to pre-pregnancy levels after delivery. Indeed, alpha diversity of the oral microbiome increases during pregnancy, and an increase of pathogenic taxa has also been reported ^36^. Methanogenic archaea are known to effectively support fermenting microorganisms by eliminating inhibiting end products (especially H_2_ and CO_2_) ^37^. Therefore, increased methanogenic archaea could be a sign of excessive growth of bacterial anaerobes at this stage, which might be associated with periodontitis. Indeed, pregnant women are more prone to dental problems ^38^.

Interpretation of signals from the mycobiome was difficult due to the large number of taxonomically unclassified reads that were received. This again highlights the lack of appropriate high-throughput methods and mycobiome databases for read annotation, especially from samples outside the gastrointestinal tract. Although the mycobiome analyses in our approach were not specifically targeting the mycobiome (e.g. adjustments in cell lysis, selection of primers or removal of host DNA could have improved results), we still obtained meaningful information: The overall mycobiome profile changed with pregnancy status, consistent with observations from previous, gut-focused studies ^39^. This change was also reflected by unknown Saccharomycetales that increased in urine samples during pregnancy. In addition, the number of samples that failed sequencing and quality controls suggested that pregnant women have lower levels of fungal microbiome components in their gastrointestinal tract.

Since the overall impact of pregnancy status on the gastrointestinal microbiome was found to be relatively small, we focused on the urogenital and oral microbiome, which showed greater restructuring. Indeed, pregnant and postpartum women are at higher risk for health problems at these body sites, such as periodontitis ^8,9^, bacterial vaginosis ^31^, or fungal infection of the vagina ^13,32,33^. Any aspects that could contribute to a rapid return to a microbiome composition typical of a healthy, nonpregnant state could help reduce these microbiome-related complications, which could also affect maternal recovery.

In agreement with other reports ^3,14,40,41^, we clearly demonstrated that the vaginal microbiome does not return to the nonpregnant state one month after delivery, but that remains substantially altered, mainly due to the decrease of *Lactobacillus*. This decline has already been reported ^14^ and it can be explained by a decrease in estrogen in the female body due to impaired ovarian activity after delivery and during lactation ^3^. Decreased abundance of lactobacilli, especially in the first six weeks postpartum may also be due to the lochial discharge, which has previously been characterized as alkaline and therefore may restrict the growth of *Lactobacillus* species ^3^. Loss of lactobacilli is also accompanied by an increase in other, often opportunistic pathogenic bacteria, leading to the predominance of vaginal community state type (CST) IV in postpartum women, which is generally associated with urogenital infections ^42,43^. During the puerperal time, women are at higher risk of vaginal postnatal infections due to the large wound area, as well as other infections such as endometritis when vaginal opportunistic bacteria enter the upper genital tract, or urinary infections ^14,44^. Probiotics treatment was already investigated in multiple other studies wo prevent or fight vaginal infections ^45–47^. It needs to be further investigated how and to what extend administration of probiotics postpartum are capable to accelerate recovery of the postpartum vaginal microbiome

Of note, mode of delivery did not appear to affect the postpartum vaginal microbiome. Since many women in this cohort who delivered vaginally had birth tears and all women who delivered by cesarean section were treated with antibiotics, we would have expected a separation of their microbiomes, but this effect was not observed.

Source tracking analysis indicated a strong exchange between the vaginal and urinary microbiome, with the overall effect of the urinary microbiome on the vaginal microbiome being greater than *vice versa*. Thus, not only microorganisms could be transferred, but also important metabolic compounds, such as lactose. The greatly increased lactose content in postpartum urine might contribute to the recovery of lactobacilli throughout the urogenital tract. Indeed, lactose added to fecal samples could specifically increase the abundance of lactobacilli in *in-vitro* experiments ^48^.

In addition to lactose, oxaloacetic acid was also increased in postpartum urine. Lactose is an important indicator of active lactation and might also be released from microbial metabolism of human milk oligosaccharides (HMOs) excreted with the urine ^13^. However, the origin of oxaloacetic acid is less clear. The increased excretion of unused oxalacetic acid could reflect an increased catabolic state during lactation ^34^. We hypothesize that the urinary metabolome retains this signature after birth as long as the woman is breastfeeding.

In general, the female body is in an exceptional state after birth and still when the woman is breastfeeding, even months after delivery. As long as lactation hormones are produced, metabolism and physiology are altered compared to a nonpregnant and non-breastfeeding state. Despite the fact that breastfeeding has numerous beneficial effects not only on the child but also on the mother, it might decelerate the return of the female body and its microbiome to the nonpregnant state ^14^. This is a very interesting research question, and longitudinal sampling of postpartum women who breastfeed and who do not breastfeed is needed to clarify this hypothesis.

### Limitations

In our dataset, only stool samples from pregnant women before delivery were compared to stool samples of nonpregnant controls, as no samples were collected after delivery. Another limitation of the samples collected is that we did not track the same individuals before and during pregnancy, but the two study groups include different individuals. This may explain that the stability observed in our dataset contrasts with literature of Nuriel-Ohayon et al ^2^ and Koren et al 2012 ^49^ describing dramatic changes in the composition of the gut microbiome from the first to the third trimester. However, the focus of this study was set on the oral and urogenital microbiome, as findings in the gastrointestinal microbiome has been specifically addressed in numerous other publications.

## Conclusion

In addition to the postpartum shift of *Lactobacillus* population in the vagina, which remains to be explored, additional work should be invested in the use of probiotics to improve postpartum maternal health. To avoid potentially long-term establishment of unhealthy CSTs and associated vaginal/urinary bacterial or fungal infections, we would suggest increasing *Lactobacillus* numbers in the vaginal niche, for example, by administering probiotics to naturally accelerate the establishment of a healthier vaginal microbiome that is better able to fight pathogens. Additionally, it would be important to explore why some women keep an optimal vaginal microbiome state after delivery characterized by a dominance of *Lactobacillus* species, as this could provide insights in the mechanisms that could be targeted for promoting an optimal vaginal microbiome postpartum. Overall, it remains clear that most of the research has focused on the health of the child, whereas the important recovery of the mother has not been fully elucidated, especially with regard to the non-intestinal microbiome.

## Materials and Methods

### Ethics statement

Research involving human material was performed in accordance with the Declaration of Helsinki and was approved by the local ethics committees (the Ethics Committee at the Medical University of Graz, Graz, Austria). The samples included in this study have been obtained under the ethics vote number 28-524 ex15/16. The study has been registered at clinicaltrials.gov (NCT04140747).

### Study design

Fifty-nine participants were enrolled in this pilot study to explore the microbial changes that occur during pregnancy, by comparing the microbiome of different body sites of a cohort of healthy pregnant women (n = 30) with the microbiome of the same body sites of a cohort of healthy nonpregnant women (n = 29). All recruited participants were healthy, were 18 years of age or older, willing to consent to all aspects of the protocol. Participants were excluded if they had any recent genitourinary infections, if they had taken any antibiotic/probiotic treatment in the last 6 months, if they smoked or took drugs, if they had HIV or HCV. Pregnant participants were also excluded if they had multiple pregnancy, membrane rupture longer than 12 h, pre-pregnancy diabetes type 1 or 2, gestational diabetes mellitus, pre-pregnancy hypertension or Preeclampsia/HELLP. Metadata from all participants are listed in available in the GitHub Repository https://github.com/CharlotteJNeumann/PerinatalMicrobiomeTRAMIC ^50^.

### Sample collection and processing

For pregnant participants, samples were collected at two time points,1-2 weeks before delivery (prepartum) and one month postpartum. The following samples were collected prepartum: stool, urine, vaginal, and oral samples, and postpartum: urine, vaginal and oral samples. For the nonpregnant women, sample collection was performed only at one time point and the samples collected were stool, urine, vaginal and oral samples. The samples were collected by the participants after they had been clearly instructed on how to collect and store the samples.

Stool and urine samples were collected using sterile collection tubes. Vaginal and oral samples were collected using FLOQSwabs (Copan, Milan, Italy). Oral samples were collected from the cheek buccal mucosa. All samples were transported to the lab on ice and stored at -80°C until further processing.

Genomic DNA was extracted from the urine, oral and vaginal specimens using the QIAamp DNA Mini Kit (QIAGEN) with some modification: before the extraction with the kit, the urine samples were centrifuged at 4400xg for 15 min, the supernatant was removed except for 500 µL that was used to resuspend the pellet. 500 µL of Lysis Buffer (sterile filtered, 20 mM Tris-HCl pH 8, 2 mM Na-EDTA, 1,2% Triton X-100) was added to the vaginal and oral swabs. To all the samples 50 µL of Lysozyme (10 mg/mL) and 6 µL of Mutanolysin (25 KU/mL) was added, followed by an incubation at 37°C for 1 h. The obtained mix was transferred to Lysing Matrix E tubes (MP Biomedicals) followed by a step of mechanical lysis at 5500 rpm for 30s two times using the MagNA Lyser Instrument (Roche, Mannheim, Germany). After the mechanical lysis the samples were centrifuged to separate the beads from the supernatant at 10000 x g for 2 min. Afterwards the DNA was extracted according to the provided instructions. The DNA was eluted in 100 µL of Elution Buffer for the vaginal and in 60 µL for the urine and oral samples.

The stool was processed using the QIAamp DNA Stool Mini Kit (QIAGEN) with some modification: around 200 mg of stool were mixed with 500 µL Inhibitex and homogenized. To the homogenized samples, 50 µL of Lysozyme (10 mg/mL) and 6 µL of Mutanolysin (25 KU/mL) was added and incubated at 37°C for 1 h. After the incubation step, 500 µL Inhibitex was added to the samples, homogenized and transferred to Lysing Matrix E tubes (MP Biomedicals) followed by a step of mechanical lysis at 6500 rpm for 30s two times using the MagNA Lyser Instrument (Roche, Mannheim, Germany). After the mechanical lysing step, the samples were incubated for 5 min at 70°C followed by a centrifugation step to separate the beads from the supernatant at 10000 x g for 3 min. The supernatant was transferred to 2 mL Eppendorf tubes and the rest of the DNA extraction was performed according to the kit protocol. The DNA was eluted in 200 mL. For all samples, the genomic DNA concentration was measured using Qubit HS. Most urine samples had a DNA concentration under the detection limit.

During DNA extraction, negative controls were also added and processed alongside.

### PCR amplification

The obtained genomic DNA was used to amplify the V4 region of the 16S rRNA gene using Illumina-tagged primers, 515FB and 806RB (Table 4). In order to identify the archaeal communities, present in the samples, a nested PCR was performed using the primer combination 344F-1041R/519F-Illu806R as described previously (Pausan et al., 2019). For the fungal communities, the ITS region was amplified using the primers ITS86F/ITS4. The PCR reactions were performed in triplicates in a final volume of 25 µL containing: TAKARA Ex Taq® buffer with MgCl2 (10 X; Takara Bio Inc., Tokyo, Japan), primers 200 nM, dNTP mix 200 µM, TAKARA Ex Taq® Polymerase 0.5 U, water (Lichrosolv®; Merck, Darmstadt, Germany), and DNA template (1-2 µL of genomic DNA). The PCR amplification conditions are listed in Table 5.

**Table 4.**
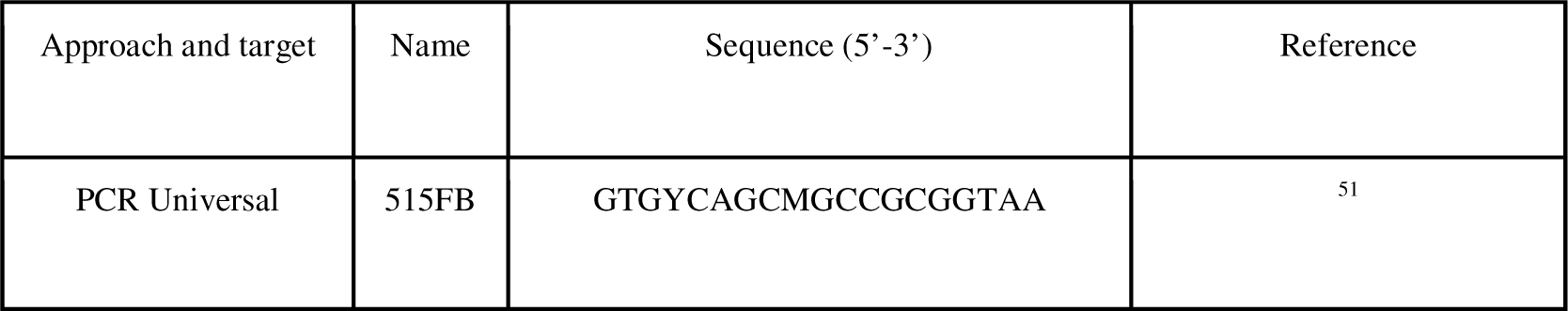
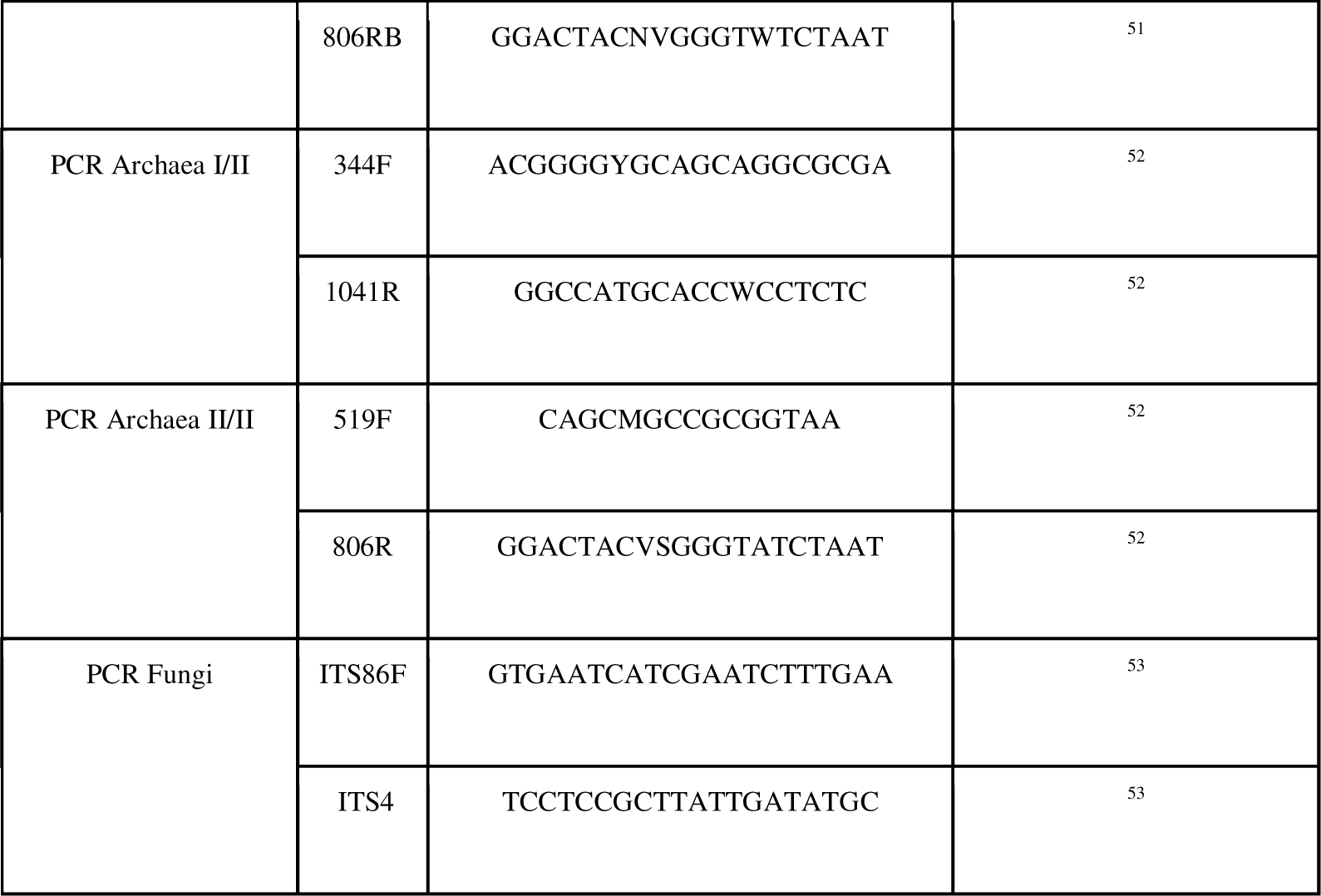
Primer pairs used for universal, archaeal and fungal PCRs.

**Table 5.** PCR settings for the primer pairs used.

| Target gene | Primer pair | Initial denaturation | Denaturation | Annealing | Elongation | Final Elongation | No. of cycles |
| --- | --- | --- | --- | --- | --- | --- | --- |
| Universal<br>(16S rRNA gene) | 515FB -<br>806RB | 3', 94°C | 45'', 94°C | 1', 50°C | 1' 30'', 72°C | 10', 72°C | 40 |
| Archaea<br>(16S rRNA gene) | 344F-1041R | 5', 95°C | 30'', 94°C | 45'', 56°C | 1', 72°C | 10', 72°C | 25 |
|  | 519F-806R | 5', 95°C | 40'', 95°C | 2', 63°C | 1', 72°C | 10', 72°C | 30 |
| Fungi<br>(ITS region) | ITS86F -<br>ITS4 | 1', 94°C | 30'', 94°C | 30'', 56°C | 30'', 68°C | 10', 68°C | 35 |

### Amplicon sequencing, bioinformatics and statistical analysis

Library preparation and sequencing of the amplicons were carried out at the Core Facility Molecular Biology, Center for Medical Research at the Medical University Graz, Austria. In brief, DNA concentrations were normalized using a SequalPrep™ normalization plate (Invitrogen), and each sample was indexed with a unique barcode sequence (8 cycles index PCR). After pooling of the indexed samples, a gel cut was carried out to purify the products of the index PCR. Sequencing was performed using the Illumina MiSeq device and MS-102-3003 MiSeq® Reagent Kit v3-600cycles (2×150 cycles). The obtained 16S rRNA gene amplicon data are available in the European Nucleotide Archive under the study accession number PRJEB65415.

The analysis of the 16S rRNA gene amplicon data was performed using QIIME2 ^54^ 2021.1-12 as described previously ^55^. Quality filtering was performed with the DADA2 algorithm ^56^ for truncation (-p-trunc-len-f 200 –p-trunc-len-r 150), and denoising to generate amplicon sequence variants (ASVs). Taxonomic classification ^57^ was based on the SILVA 138 database ^58^ and the obtained feature table and taxonomy file were used for further analysis (Supplementary Tab. 3). Contaminating ASVs were determined and removed by decontam v 1.13 ^59^ in R ^60^, running *iscontaminant* in *prevalence* mode with a threshold of 0.5. After this step, positive and negative controls were removed from the datasets. Additionally, ASVs classified as chloroplast and mitochondria were removed.

For normalization, different approaches were used for the three bacterial, fungal and archaeal datasets based on their composition. SRS (scaling with ranked subsampling) normalization was run in QIIME2 ^54^ with c_min_ = 1,500 for the bacterial dataset and with c_min_ = 100. The archaeal dataset was normalized with TSS (total sum normalization). The number of samples that were analyzed and the number of how many samples were kept after normalization is listed in Suppl. Fig. 1.

Differentially abundant taxa were defined by q2-aldex2 ^61–63^ in QIIME2 ^54^. To display those taxa in boxplots in R (packages: ggplot2 ^64^, dplyr ^65^, reshape ^66^), the data of relative abundance were first CLR transformed in R ^60^.

Source tracking was performed with Sourcetracker2 ^67^ (standard settings) with rarefaction done at 1500 sequences. Data were then visualized by Rawgraphs ^68^, based on the median of each sample set.

Sequences classified as *Lactobacillus* genus were further classified to species level to allow the clustering of the vaginal microbiome into community state types (CSTs). The classification was performed by classification through EzBioCloud ^69^. The vaginal community state types (CSTs) were classified using the VALENCIA (VAginaL community state typE Nearest CentroId clAssifier) classifier ^30^ as described in the GitHub tutorial ^70^. The biom table used to run the classifier VALENCIA as well as the obtained output can be found in the Github Repository.

A subset of 87 urine samples from all three groups were analyzed in house with untargeted NMR (nuclear magnetic resonance spectroscopy) for several metabolites as described previously ^71^. In short, methanol water was added to the samples, cells were lysed, lyophilized, and mixed with NMR buffer. NMR was performed on an AVANCETM Neo Bruker Ultrashield 600MHz spectrometer equipped with a TXI probe head at 310 K and processed as described elsewhere ^72^. NMR data were analyzed using MetaboAnalyst ^73^, following the protocol for paired samples (comparison mpre to mpost; log transformation).

Significantly differentially abundant metabolites were correlated with CLR transformed relative abundance of bacterial genera in R ^60^ and plotted in heatmaps with ggplot2 ^64^.

The circle packing plot about archaeal occurrence was created with rawgraphs.io ^68^. All figures were adjusted in inkscape v 1.1 (URL: https://inkscape.org/en/RRID:SCR_014479).

All data tables are available in our Github Repository https://github.com/CharlotteJNeumann/PerinatalMicrobiomeTRAMIC ^50^.

An overview of the available data is displayed in two figures: Suppl. Fig. 1 is following the STORM guideline and was created with drawio.com (URL: https://drawio.com). Suppl. Fig. 2 displays the data available per sample and individual.

## Supplementary FIGURES COLLECTION

**Suppl. Fig. 1:**
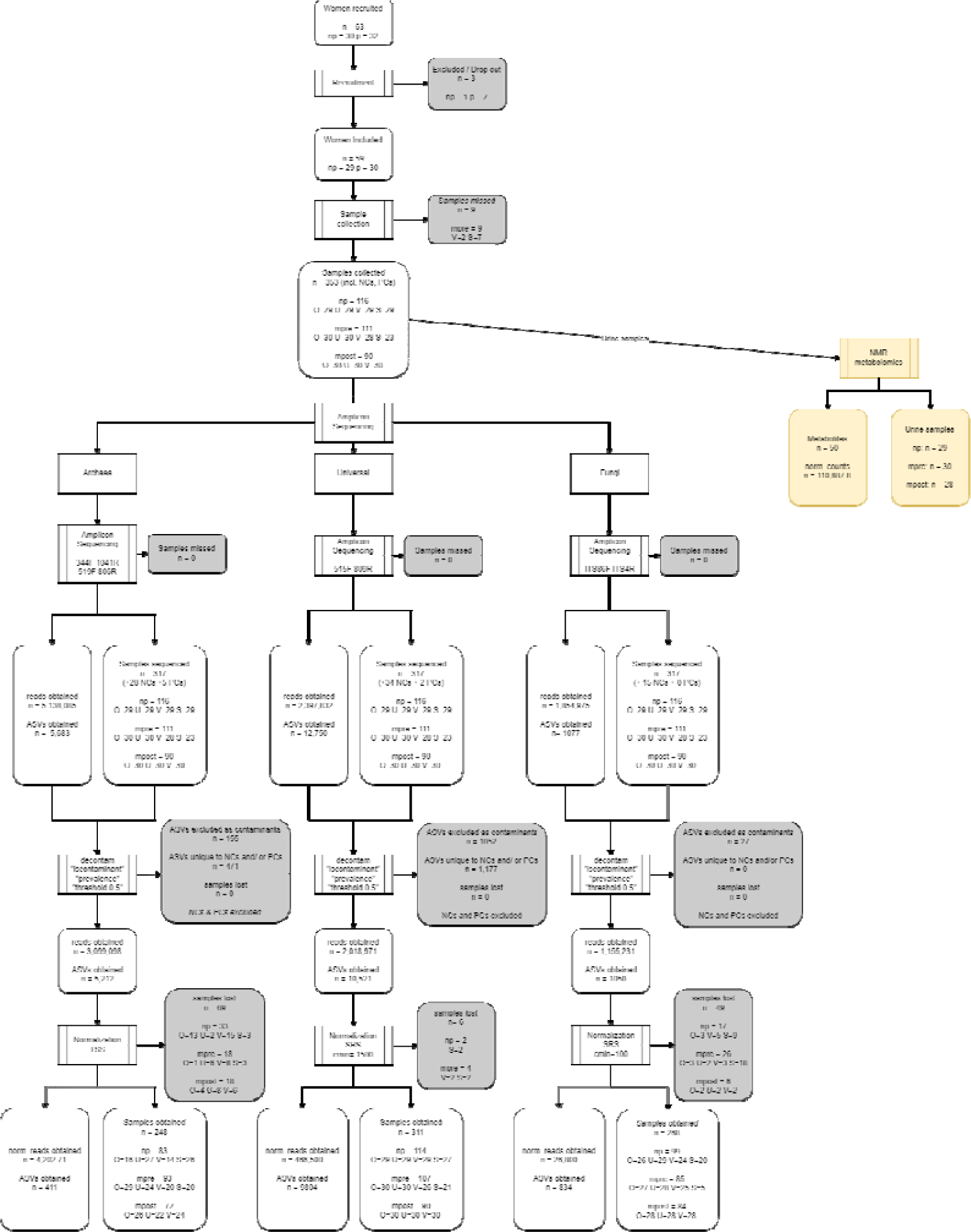
flowchart according to the STORM pipeline, showing sample and read numbers for data analyses steps.

**Suppl. Fig. 2:**
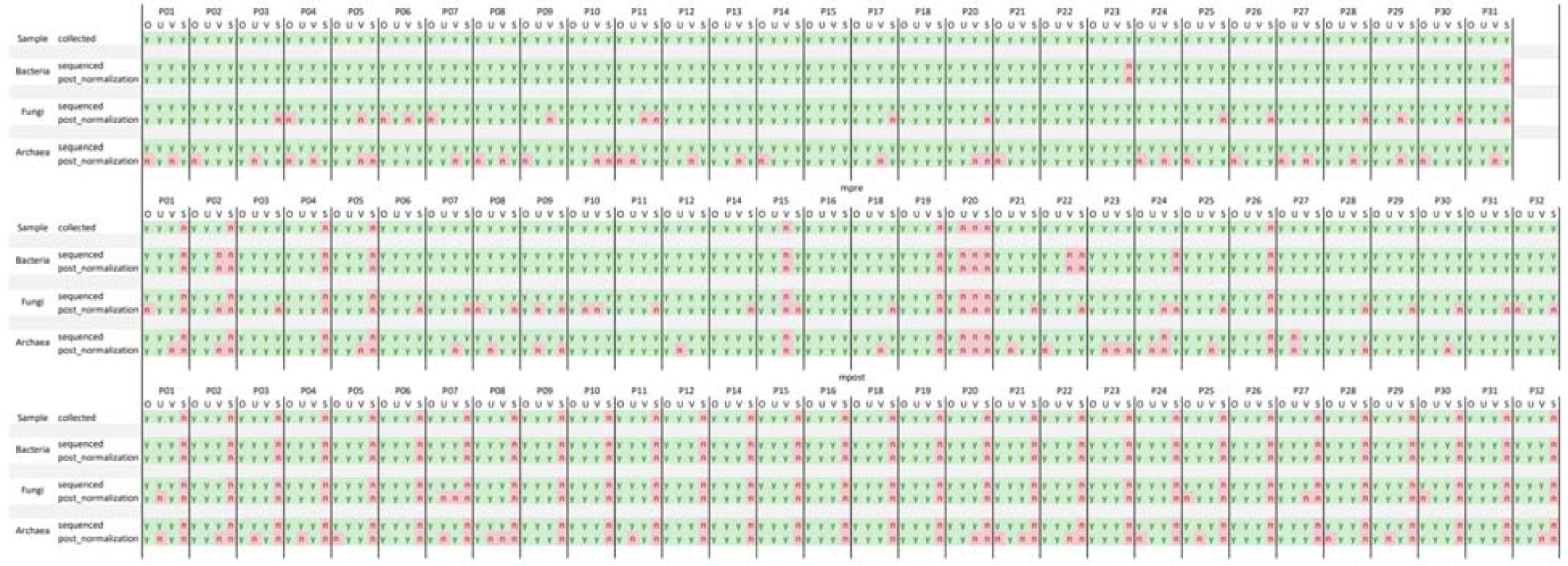
Sample Overview for nonpregnant (up), mpre (middle) and mpost (down) for single individuals; it is shown if samples are available (y, green) or not (n, red) at the steps of sample collection, sequencing and post-normalization, each for bacteria, fungi and archaea.

**Suppl. Fig. 3:**
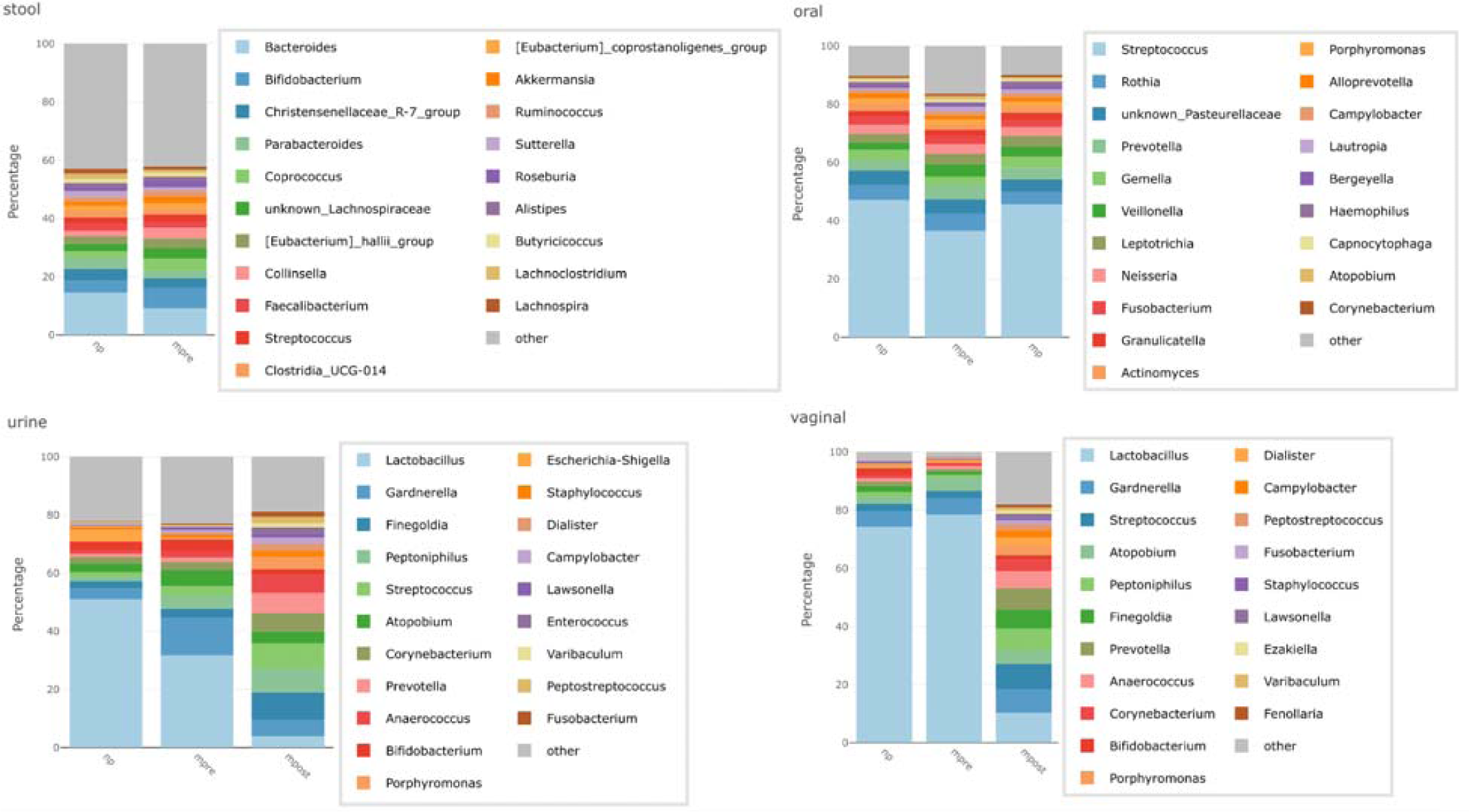
stacked bar plots showing relative abundances of the top 20 most abundant bacterial genera depicted per group for stool, oral, urine and vaginal samples.

**Suppl. Fig. 4:**
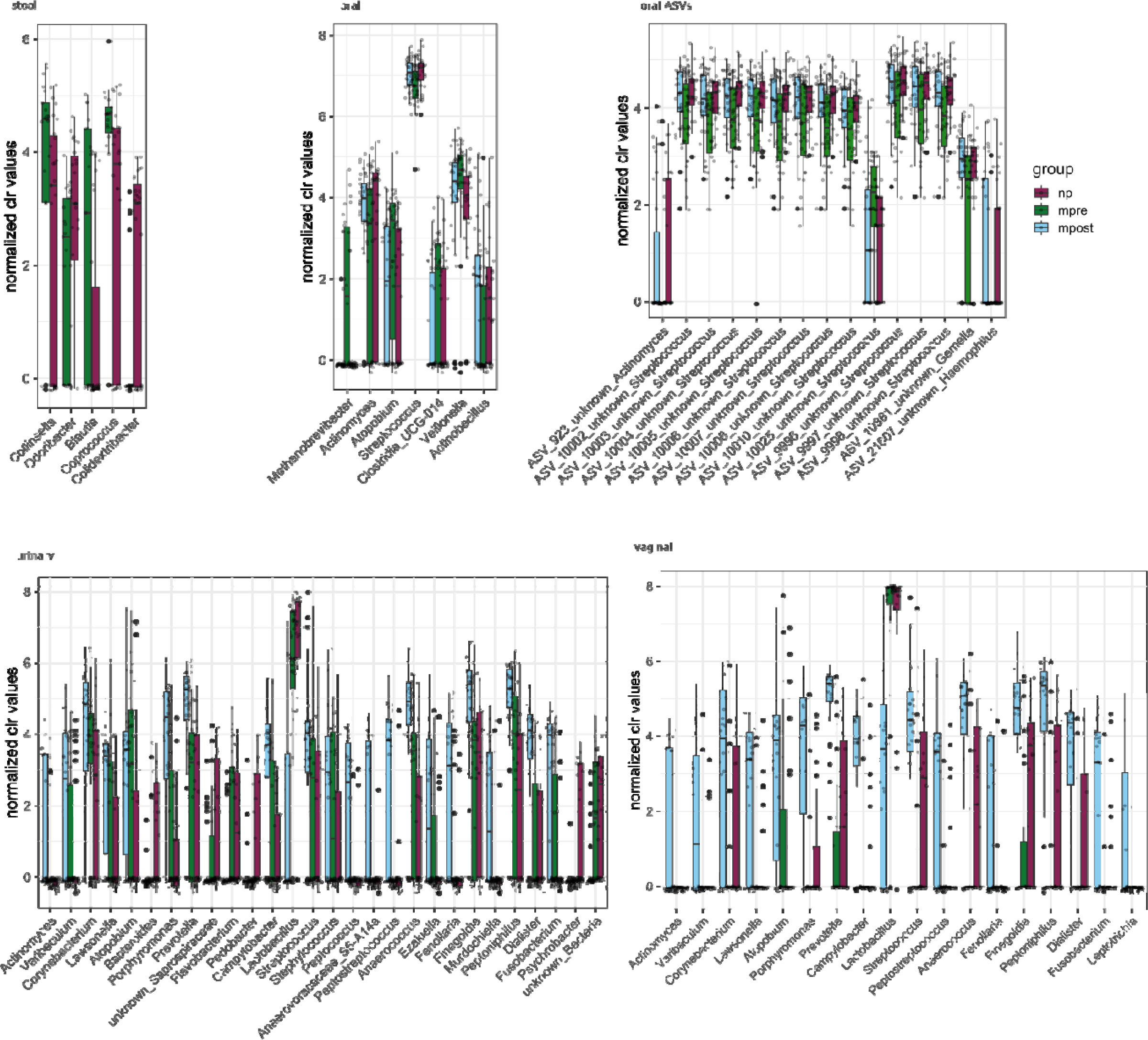
Differentially abundance boxplots of CLR transformed values on bacterial genera (Aldex2) for stool, oral, urine and vaginal samples. For oral, additionally ASVs are also depicted.

**Suppl. Fig. 5:**
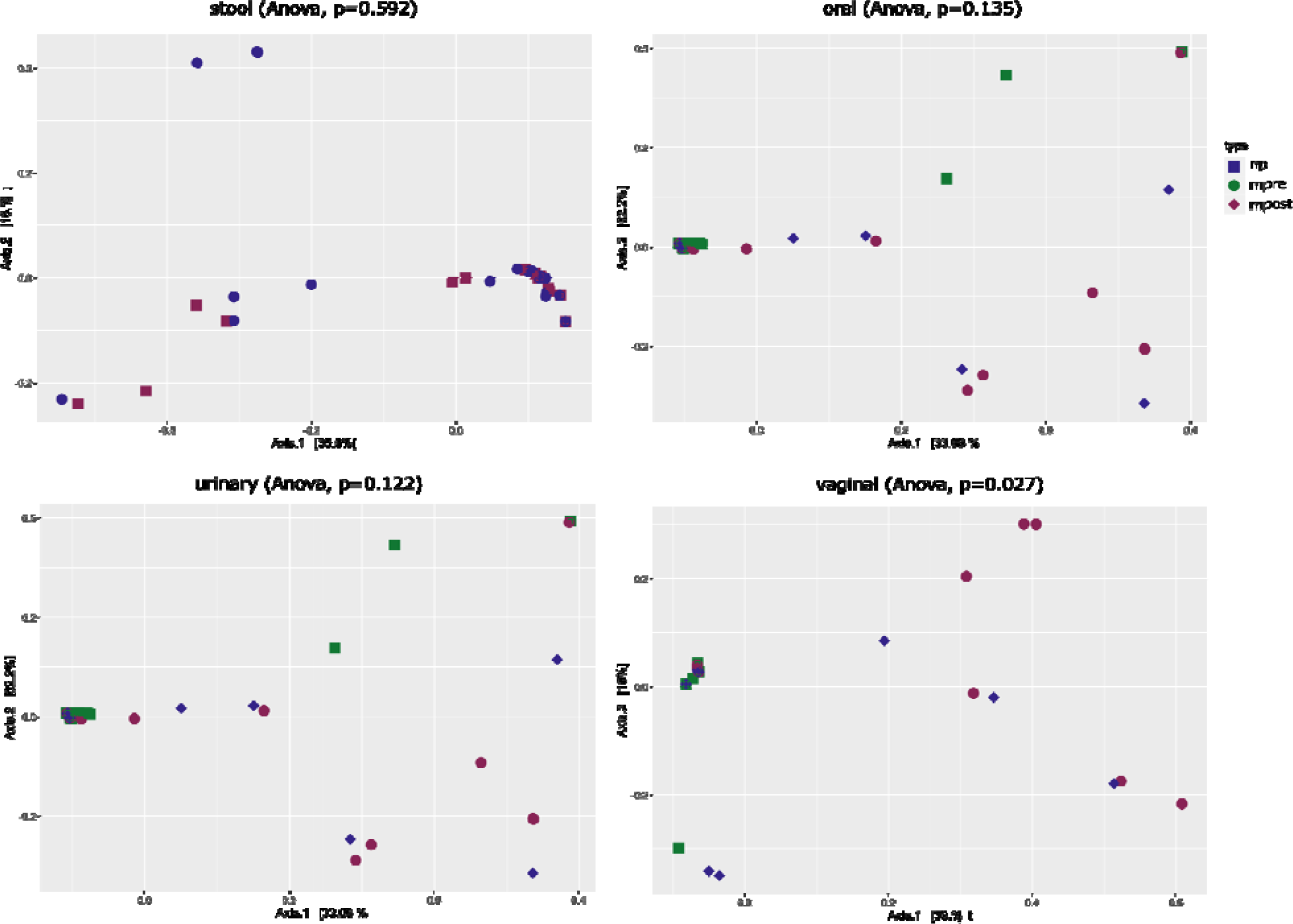
Principal Coordinate Analysis (PCoA) for groups (np, mpre, mpost) for archaeal ASVs with Unweighted Unifrac as distance Matrix and Anova p-values.

**Suppl. Fig. 6:**
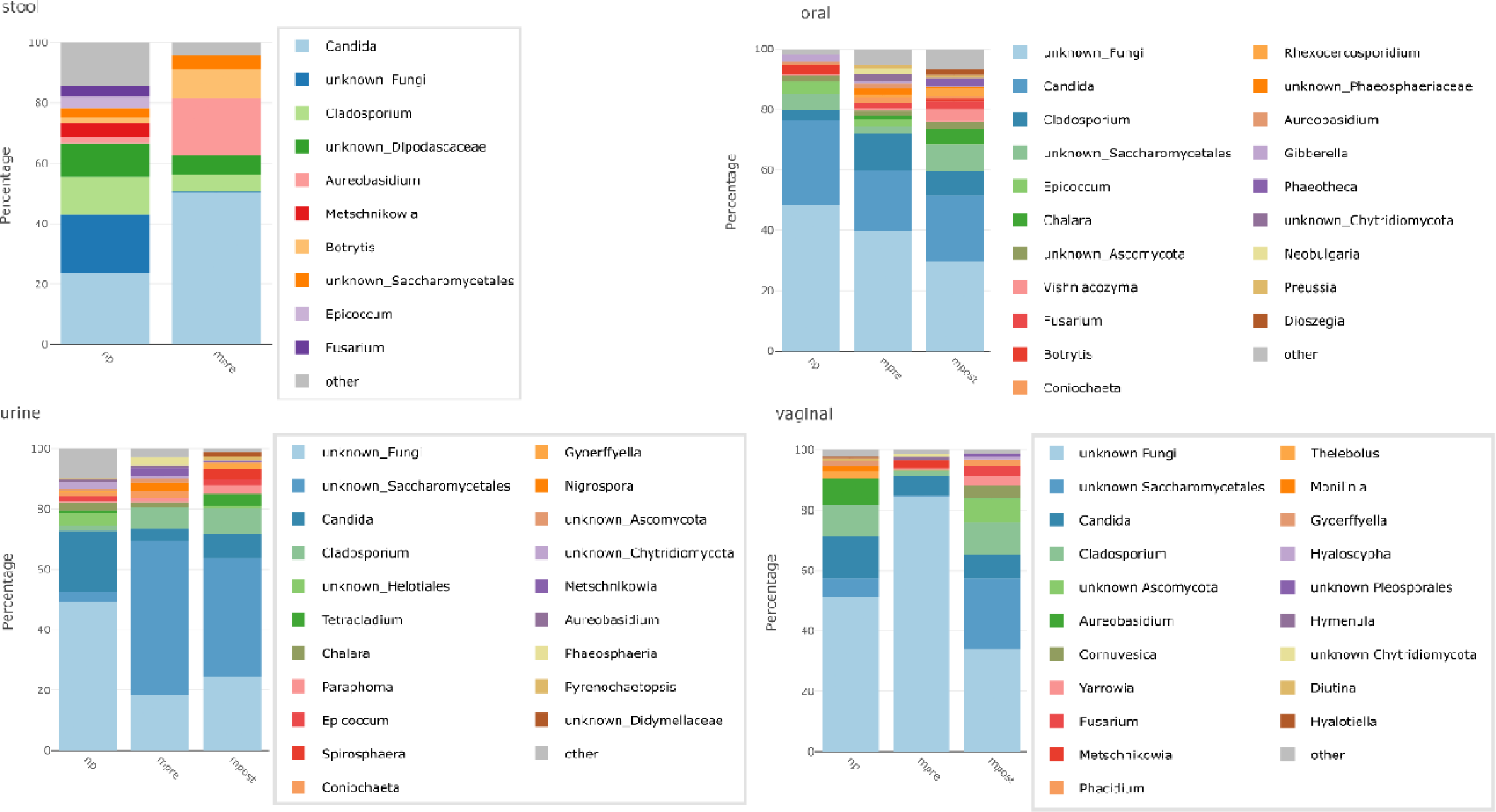
stacked bar plots showing relative abundances of the top 10 or 20 most abundant fungal genera depicted per group for stool, oral, urine and vaginal samples.

**Suppl. Fig. 7:**
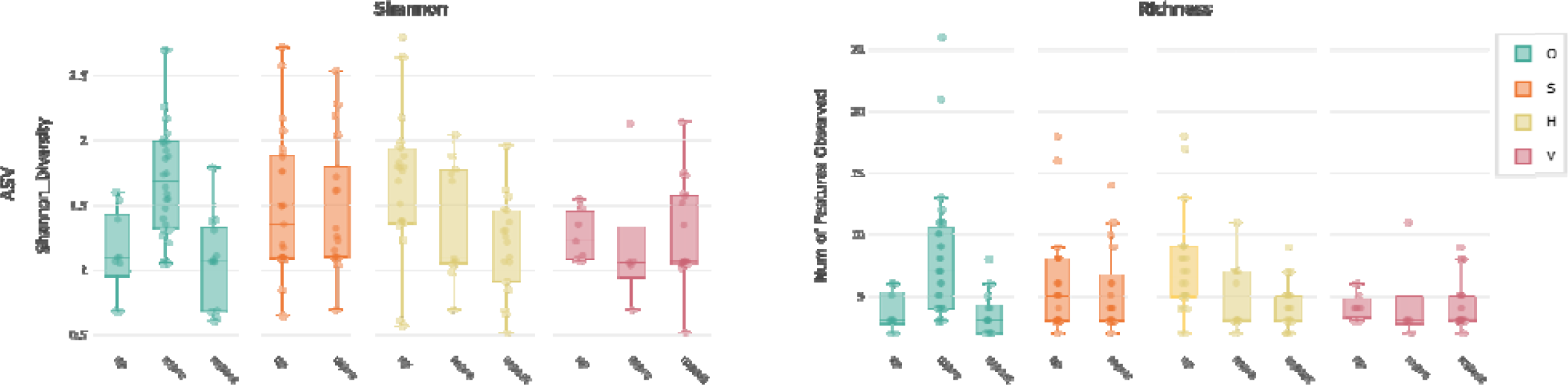
alpha diversity of the archaeal microbiome on ASV level, archaea nested PCR approach: Shannon diversity and richness, depicted per body site and split by group *p* < 0.05*, *p* < 0.005**, *p* < 0.001***

**Suppl. Fig 8:**
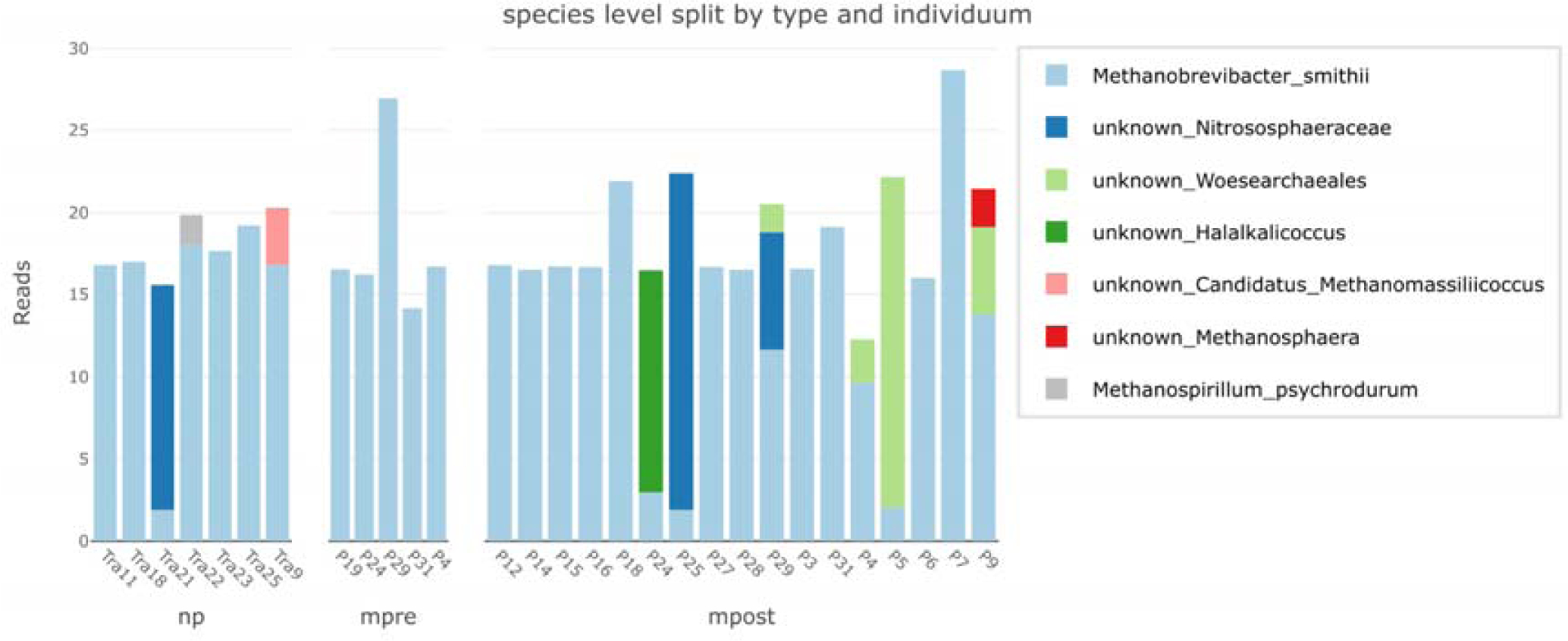
stacked bar plots showing relative abundances of the topmost abundant archaeal genera depicted per group for stool, oral, urine and vaginal samples.

**Suppl. Fig. 9:**
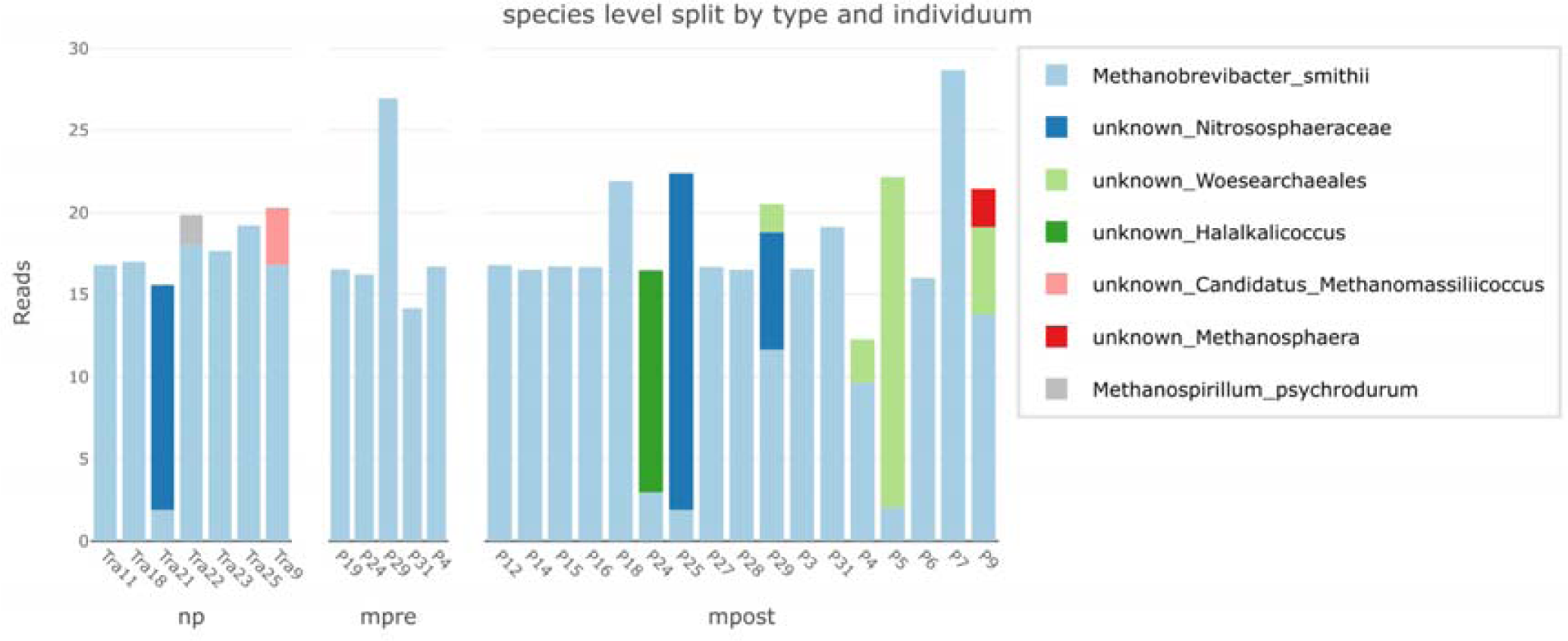
Occurrence (reads, archaea nested PCR approach) of archaeal species in the vaginal microbiome, split by group per individual.

**Suppl. Fig. 10:**
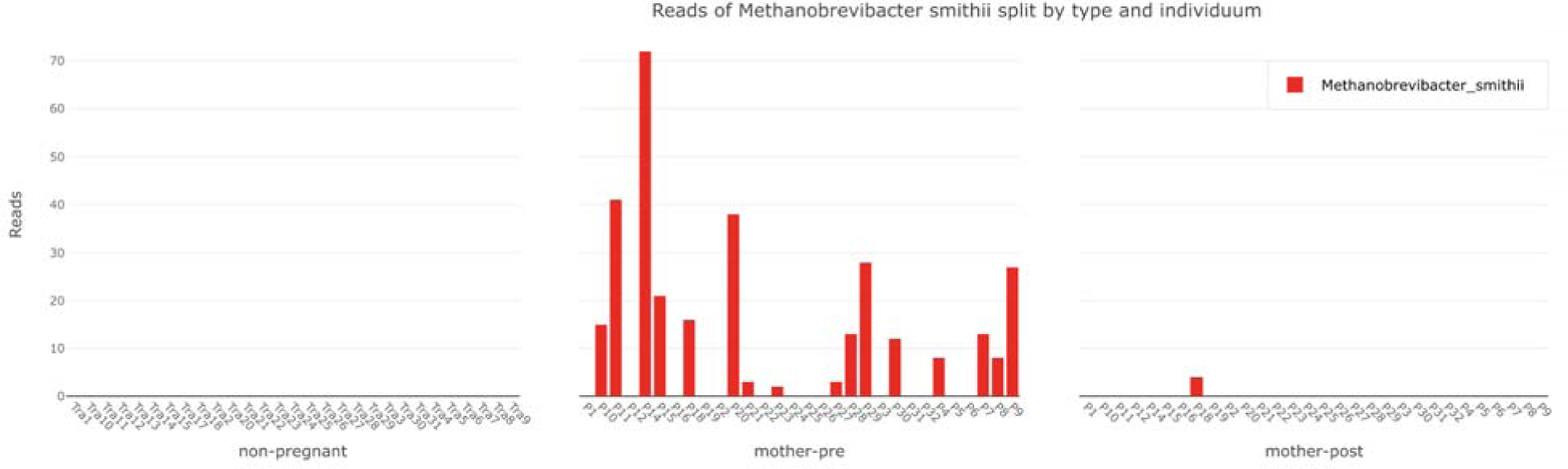
Occurrence (reads, Caporaso Approach) of *M. smithii* in the oral microbiome, split by ty e per individual.

**Suppl. Fig. 11:**
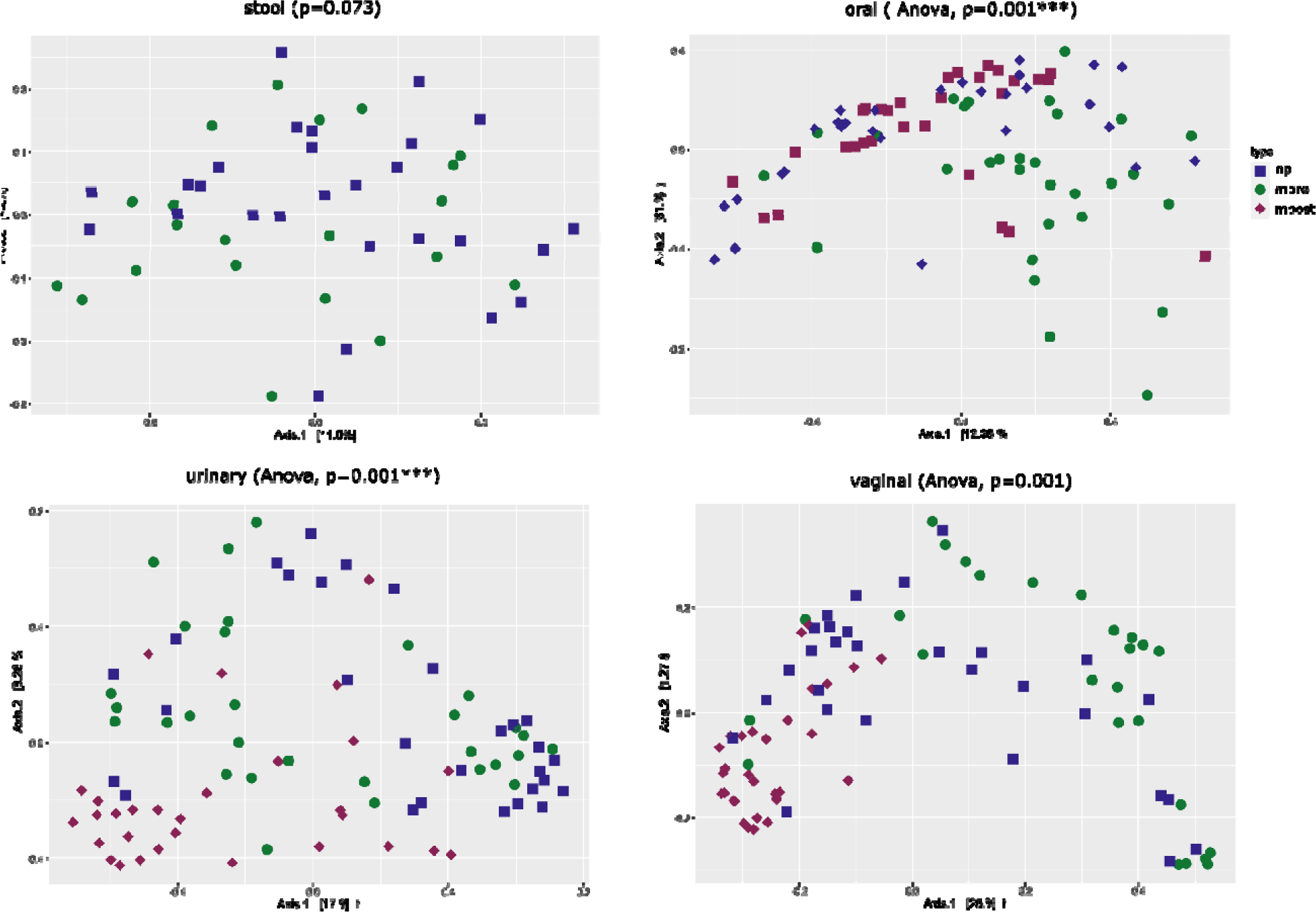
Principal Coordinate Analysis (PCoA) for groups (np, mpre, mpost) for bacterial ASVs with Unweig Unifrac as distance Matrix and Anova p-values.

**Suppl. Fig. 12:**
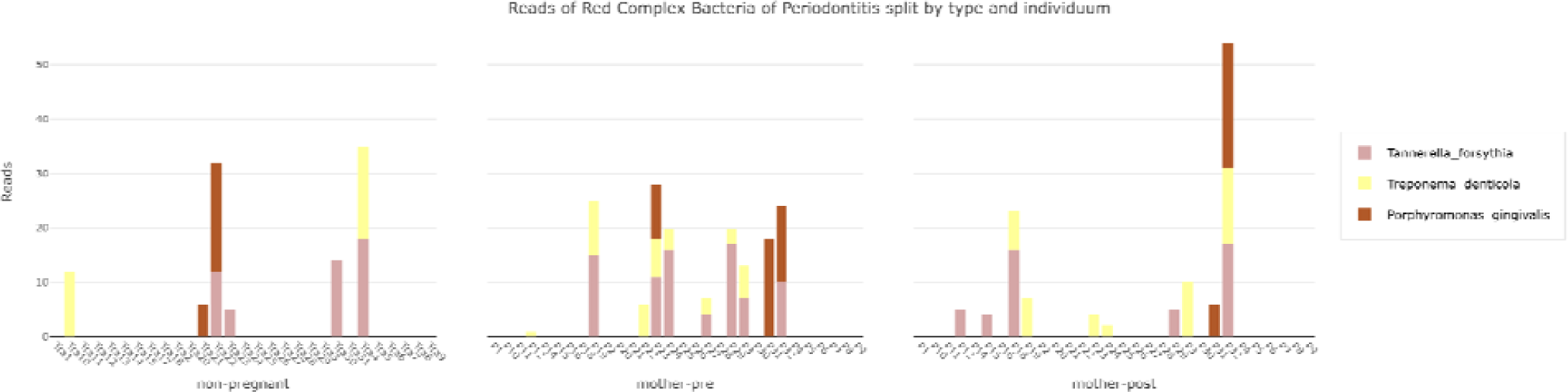
different figures on red complex bacteria in the oral microbiome.

**Suppl. Fig. 13:**
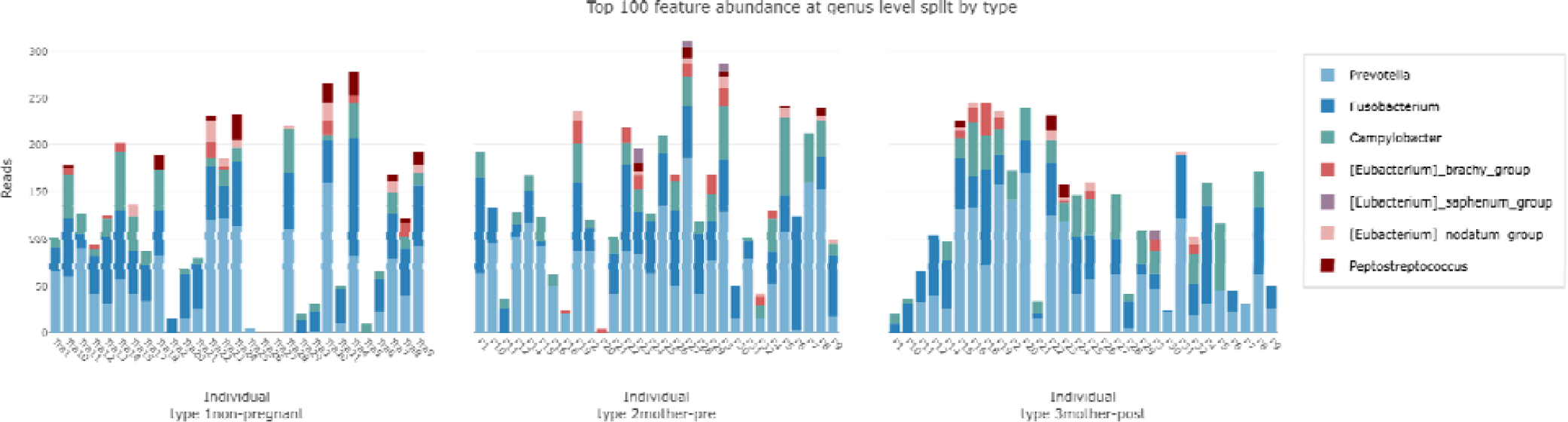
orange complex bacteria in the oral microbiome.

**Suppl. Fig. 14:**
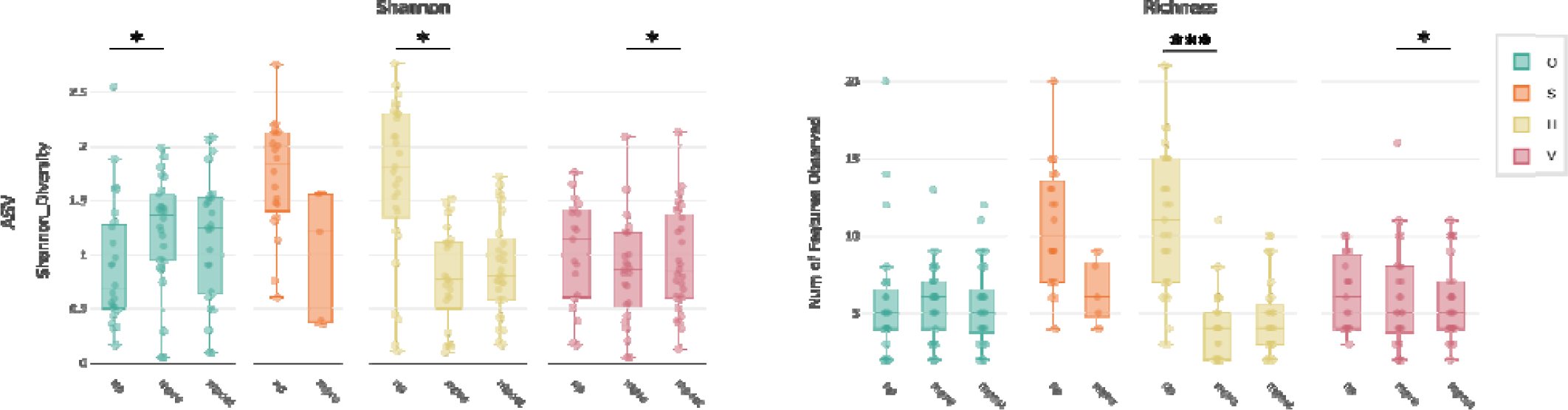
alpha diversity of the fungal microbiome on ASV level: Shannon diversity and richness, depicted per body site and split by group *p* < 0.05*, *p* < 0.005**, *p* < 0.001***

**Suppl. Fig. 15:**
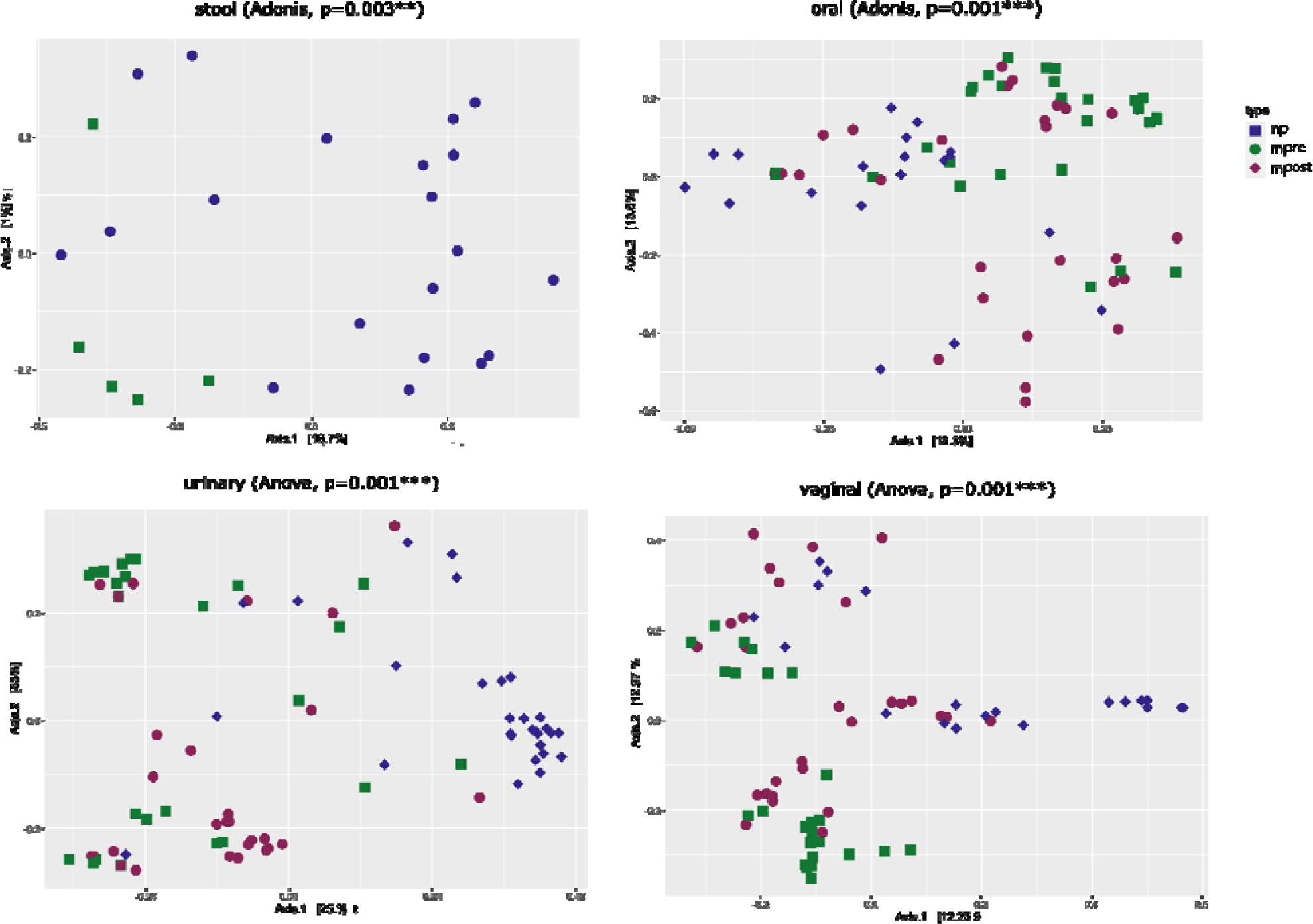
Principle Coordinate Analysis (PCoA) for groups (np, mpre, mpost) for fungal ASVs with Unweighted Unifrac as distance Matrix and Anova p-values.

**Suppl. Fig. 16:**
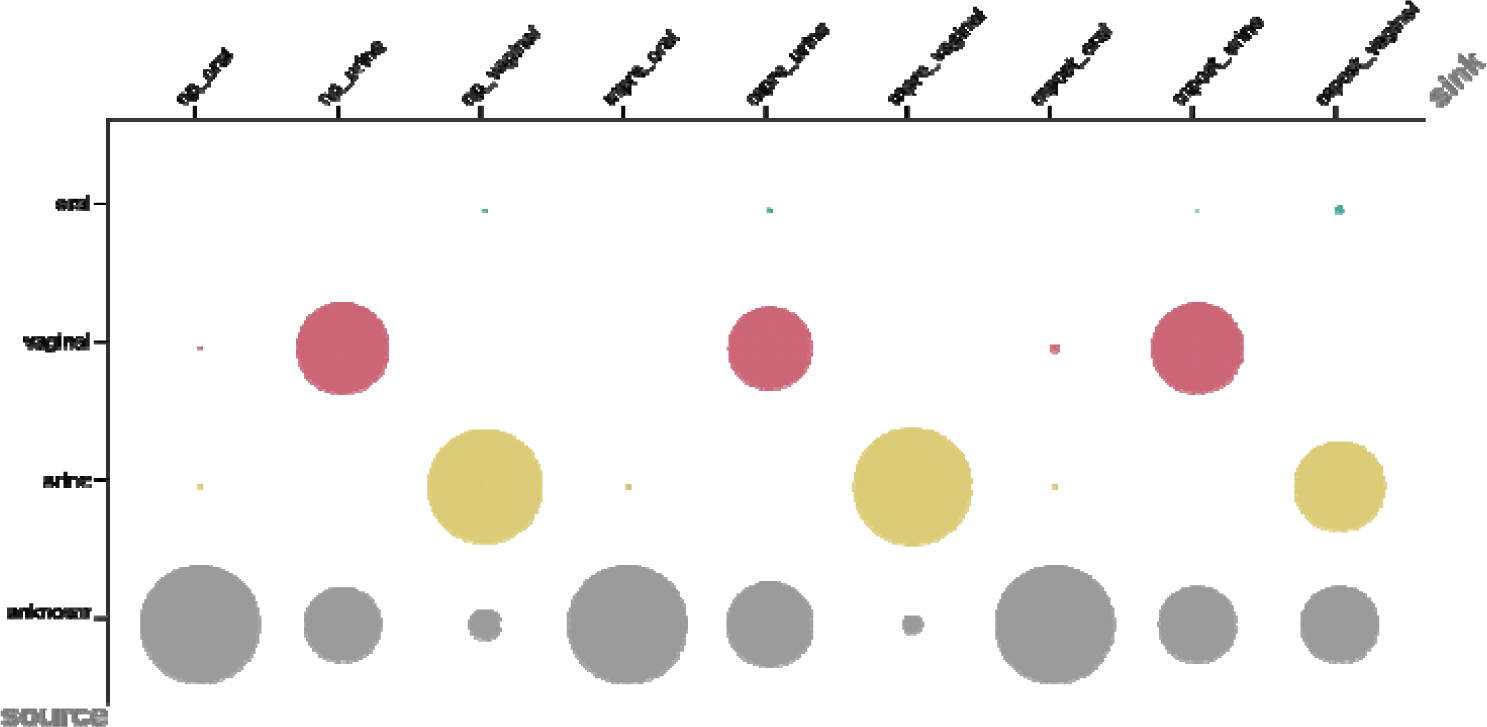
Source tracking; comparison of oral, vaginal urine samples with respect to their contribution from oral, urine, vaginal and unknown sources; dot sizes represent the proportion of contribution (numbers given in the text).

**Suppl. Fig. 17:**
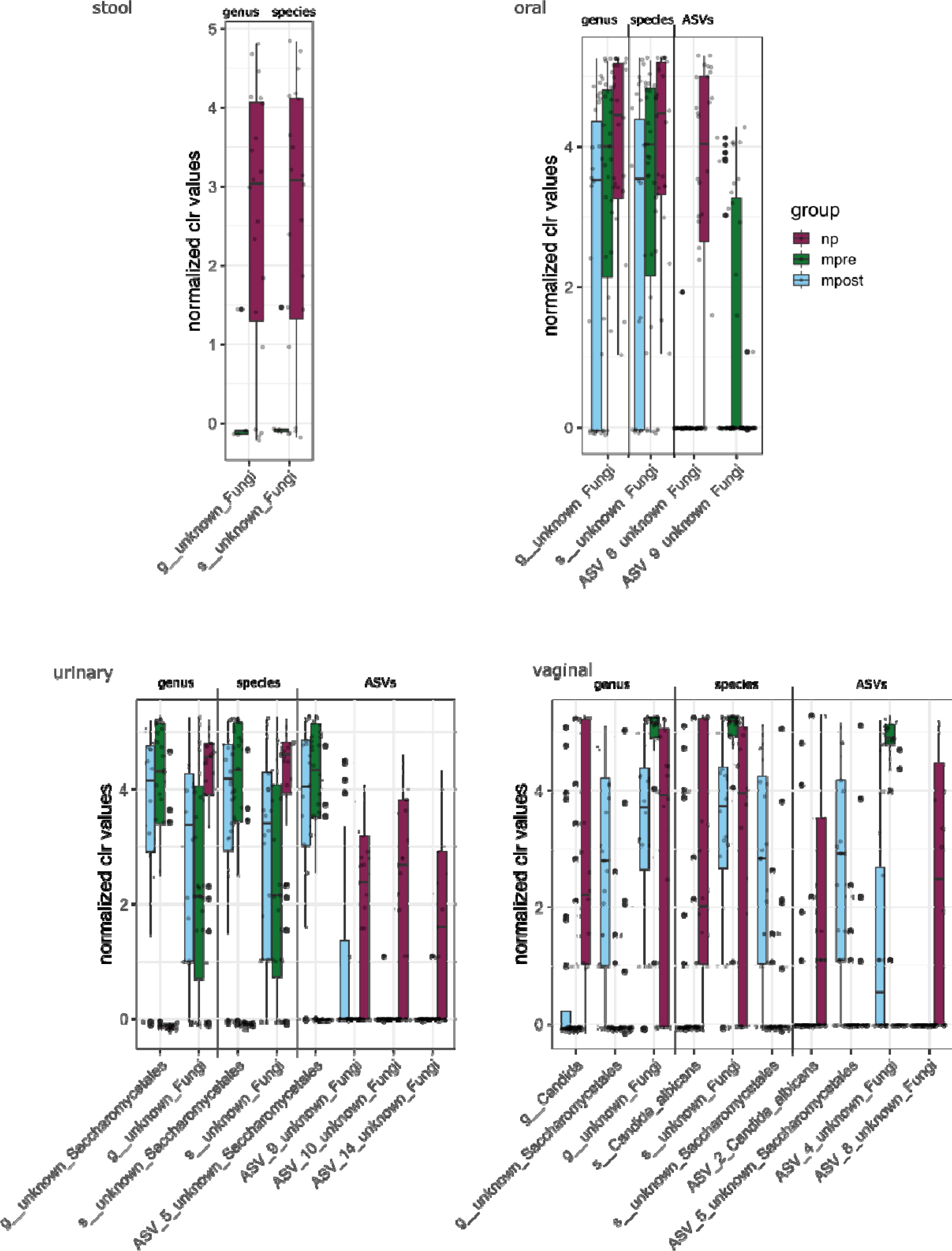
Differentially abundance boxplots of CLR transformed values on fungal genera (Aldex2) for stool, oral, urine and vaginal samples.

**Suppl. Fig. 18:**
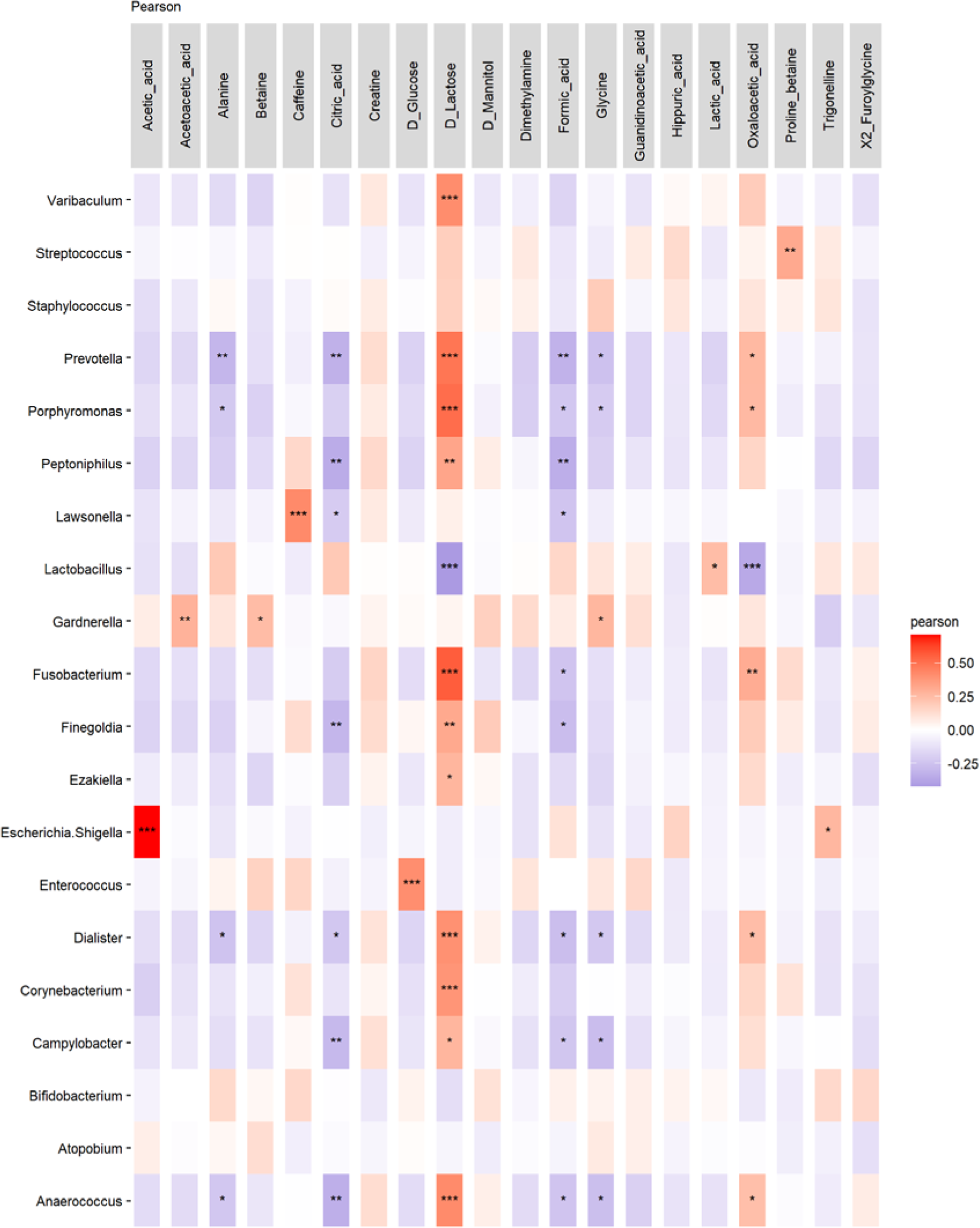
heat map of metabolites that are significantly differentially abundant between mpre and mpost (see Volcano plot figure 4A) in a pearson-correlation analysis with the taxonomic data.

## Acknowledgements

The authors thank the Medical University of Graz for the computational resources of the MedBioNode and the Life Science Compute Cluster (LiSC) operated by the Computational Systems Biology group at the University of Vienna. We thank the Medical University of Graz ZMF Galaxy Team: Core Facility Computational Bioanalytics, Medical University of Graz, funded by the Austrian Federal Ministry of Education, Science and Research, Hochschulraum-Strukturmittel 2016 grant as part of BioTechMed Gral 2016 grant as part of BioTechMed Graz.

We thank Tobias Madl and his research team at Medical University of Graz for NMR Metabolomics analysis and the Department of Obstetrics and Gynecology of Meduni Graz for sample collection. A special thanks goes to the participants of this study for providing samples and information.

## Funding details

This research was funded by the Austrian Science Fund (FWF), under project number KLI 784. The study was financially supported by the City of Graz (to MRP) and the Austrian Commission for UNESCO and L’ORÉAL with the L’OREAL Fellowship for Women in Science (to M.-R.P.). M-R.P. was trained in the Doctoral Program MolMed, and C.N. was trained in the PhD Program DP-iDP at the Medical University of Graz.

## Disclosure statement

The authors report there are no competing interests to declare.

Data are available in the European Nucleotide Archive under the study accession number PRJEB65415. Data and Supplemental material are deposited in our GitHub Repository https://github.com/CharlotteJNeumann/PerinatalMicrobiomeTRAMIC ^50^ DOI: 674556830

## Contributions

Conceptualization: M.-R.P., E-C.W., V.K-K., E.J-K., C.M-E. Methodology: C.J.N., M.-R.P., A.M., C.M.-E. Formal Analysis: C.J.N., M.-R.P., C.M.-E. Investigation: C.J.N., M-R.P., V.H., B.A., P.W. Writing original draft: C.J.N., M.-R.P., C.M.-E. Writing review and editing: C.J.N., M.-R.P., V.H., E-C.W., V.K-K., B.A., P.W., A.M., E.J-K., C.M-E. Visualization: C.J.N., M-R.P.; C.M-E. Supervision: C.M-E., E.J-K. Project Administration: E-C.W., V.K-K., B.A., P.W. Funding Acquisition: C.J.N., M-R.P., C.M.-E., E.J-K.

## Notes

https://github.com/CharlotteJNeumann/PerinatalMicrobiomeTRAMIC

### Competing Interest Statement

The authors have declared no competing interest.

