## Supplementary Figure 2 for "The Dynamics of the Female Microbiome: Unveiling Abrupt Changes of Microbial Domains across Body Sites from Preconception to Perinatal Phase"

| | | P01 | | | | P02 | | | | P03 | | | | P04 | | | | P05 | | | | P06 | | | | P07 | | | | P08 | | | |
|---|---|---|---|---|---|---|---|---|---|---|---|---|---|---|---|---|---|---|---|---|---|---|---|---|---|---|---|---|---|---|---|---|---|
| | | O | U | V | S | O | U | V | S | O | U | V | S | O | U | V | S | O | U | V | S | O | U | V | S | O | U | V | S | O | U | V | S |
| Sample | collected | y | y | y | y | y | y | y | y | y | y | y | y | y | y | y | y | y | y | y | y | y | y | y | y | y | y | y | y | y | y | y | y |
| Bacteria | sequenced | y | y | y | y | y | y | y | y | y | y | y | y | y | y | y | y | y | y | y | y | y | y | y | y | y | y | y | y | y | y | y | y |
| | post_normalization | y | y | y | y | y | y | y | y | y | y | y | y | y | y | y | y | y | y | y | y | y | y | y | y | y | y | y | y | y | y | y | y |
| Fungi | sequenced | y | y | y | y | y | y | y | y | y | y | y | y | y | y | y | y | y | y | y | y | y | y | y | y | y | y | y | y | y | y | y | y |
| | post_normalization | y | y | y | y | y | y | y | y | y | y | y | n | n | y | y | y | y | y | n | y | n | y | n | y | n | y | y | y | y | y | y | y |
| Archaea | sequenced | y | y | y | y | y | y | y | y | y | y | y | y | y | y | y | y | y | y | y | y | y | y | y | y | y | y | y | y | y | y | y | y |
| | post_normalization | n | y | n | y | n | y | y | y | y | n | y | y | n | y | n | y | y | y | n | n | y | y | y | y | y | y | n | y | n | y | n | y |

| | | P09 | | | | P10 | | | | P11 | | | | P12 | | | | P13 | | | | P14 | | | | P15 | | | | P17 | | | |
|---|---|---|---|---|---|---|---|---|---|---|---|---|---|---|---|---|---|---|---|---|---|---|---|---|---|---|---|---|---|---|---|---|---|
| | | O | U | V | S | O | U | V | S | O | U | V | S | O | U | V | S | O | U | V | S | O | U | V | S | O | U | V | S | O | U | V | S |
| Sample | collected | y | y | y | y | y | y | y | y | y | y | y | y | y | y | y | y | y | y | y | y | y | y | y | y | y | y | y | y | y | y | y | y |
| Bacteria | sequenced | y | y | y | y | y | y | y | y | y | y | y | y | y | y | y | y | y | y | y | y | y | y | y | y | y | y | y | y | y | y | y | y |
| | post_normalization | y | y | y | y | y | y | y | y | y | y | y | y | y | y | y | y | y | y | y | y | y | y | y | y | y | y | y | y | y | y | y | y |
| Fungi | sequenced | y | y | y | y | y | y | y | y | y | y | y | y | y | y | y | y | y | y | y | y | y | y | y | y | y | y | y | y | y | y | y | y |
| | post_normalization | y | y | n | y | y | y | y | y | y | y | n | n | y | y | y | y | y | y | y | y | y | y | y | y | y | y | y | y | y | y | y | n |
| Archaea | sequenced | y | y | y | y | y | y | y | y | y | y | y | y | y | y | y | y | y | y | y | y | y | y | y | y | y | y | y | y | y | y | y | y |
| | post_normalization | n | y | y | y | y | y | n | n | n | n | y | y | y | y | n | y | y | y | n | y | n | y | y | y | y | y | y | y | y | y | n | y |

| | | P18 | | | | P20 | | | | P21 | | | | P22 | | | | P23 | | | | P24 | | | | P25 | | | | P26 | | | |
|---|---|---|---|---|---|---|---|---|---|---|---|---|---|---|---|---|---|---|---|---|---|---|---|---|---|---|---|---|---|---|---|---|---|
| | | O | U | V | S | O | U | V | S | O | U | V | S | O | U | V | S | O | U | V | S | O | U | V | S | O | U | V | S | O | U | V | S |
| Sample | collected | y | y | y | y | y | y | y | y | y | y | y | y | y | y | y | y | y | y | y | y | y | y | y | y | y | y | y | y | y | y | y | y |
| Bacteria | sequenced | y | y | y | y | y | y | y | y | y | y | y | y | y | y | y | y | y | y | y | n | y | y | y | y | y | y | y | y | y | y | y | y |
| | post_normalization | y | y | y | y | y | y | y | y | y | y | y | y | y | y | y | y | y | y | y | n | y | y | y | y | y | y | y | y | y | y | y | y |
| Fungi | sequenced | y | y | y | y | y | y | y | y | y | y | y | y | y | y | y | y | y | y | y | y | y | y | y | y | y | y | y | y | y | y | y | y |
| | post_normalization | y | y | y | y | y | y | y | n | y | y | y | y | y | y | y | y | y | y | y | y | y | y | y | y | y | y | y | n | y | y | y | n |
| Archaea | sequenced | y | y | y | y | y | y | y | y | y | y | y | y | y | y | y | y | y | y | y | y | y | y | y | y | y | y | y | y | y | y | y | y |
| | post_normalization | y | y | y | y | y | y | n | n | n | y | y | y | y | y | y | y | y | y | y | y | n | y | n | y | n | y | y | y | n | y | y | y |

| | | P27 | | | | P28 | | | | P29 | | | | P30 | | | | P31 | | | |
|---|---|---|---|---|---|---|---|---|---|---|---|---|---|---|---|---|---|---|---|---|---|
| | | O | U | V | S | O | U | V | S | O | U | V | S | O | U | V | S | O | U | V | S |
| Sample | collected | y | y | y | y | y | y | y | y | y | y | y | y | y | y | y | y | y | y | y | y |
| Bacteria | sequenced | y | y | y | y | y | y | y | y | y | y | y | y | y | y | y | y | y | y | y | n |
| | post_normalization | y | y | y | y | y | y | y | y | y | y | y | y | y | y | y | y | y | y | y | n |
| Fungi | sequenced | y | y | y | y | y | y | y | y | y | y | y | y | y | y | y | y | y | y | y | y |
| | post_normalization | y | y | y | y | y | y | y | n | y | y | n | y | y | y | y | n | y | y | y | n |
| Archaea | sequenced | y | y | y | y | y | y | y | y | y | y | y | y | y | y | y | y | y | y | y | y |
| | post_normalization | n | y | n | y | y | y | n | y | y | y | n | y | n | y | y | y | y | y | n | y |

mpre

| | | P01 | | | | P02 | | | | P03 | | | | P04 | | | | P05 | | | | P06 | | | | P07 | | | | P08 | | | |
|---|---|---|---|---|---|---|---|---|---|---|---|---|---|---|---|---|---|---|---|---|---|---|---|---|---|---|---|---|---|---|---|---|---|
| | | O | U | V | S | O | U | V | S | O | U | V | S | O | U | V | S | O | U | V | S | O | U | V | S | O | U | V | S | O | U | V | S |
| Sample | collected | y | y | y | n | y | y | y | n | y | y | y | y | y | y | y | n | y | y | y | n | y | y | y | y | y | y | y | y | y | y | y | y |
| Bacteria | sequenced | y | y | y | n | y | y | n | n | y | y | y | y | y | y | y | n | y | y | y | n | y | y | y | y | y | y | y | y | y | y | y | y |
| | post_normalization | y | y | y | n | y | y | n | n | y | y | y | y | y | y | y | n | y | y | y | n | y | y | y | y | y | y | y | y | y | y | y | y |
| Fungi | sequenced | y | y | y | n | y | y | y | n | y | y | y | y | y | y | y | n | y | y | y | n | y | y | y | y | y | y | y | y | y | y | y | y |
| | post_normalization | n | y | y | n | y | y | n | n | y | y | y | n | y | y | y | n | y | y | y | n | y | y | y | n | y | y | y | n | n | y | y | n |
| Archaea | sequenced | y | y | y | n | y | y | y | n | y | y | y | y | y | y | y | n | y | y | y | n | y | y | y | y | y | y | y | y | y | y | y | y |
| | post_normalization | y | y | n | n | y | y | n | n | y | y | y | y | y | y | y | n | y | y | n | n | y | y | y | y | y | y | n | y | y | n | y | y |

| | | P09 | | | | P10 | | | | P11 | | | | P12 | | | | P14 | | | | P15 | | | | P16 | | | | P18 | | | |
|---|---|---|---|---|---|---|---|---|---|---|---|---|---|---|---|---|---|---|---|---|---|---|---|---|---|---|---|---|---|---|---|---|---|
| | | O | U | V | S | O | U | V | S | O | U | V | S | O | U | V | S | O | U | V | S | O | U | V | S | O | U | V | S | O | U | V | S |
| Sample | collected | y | y | y | y | y | y | y | y | y | y | y | y | y | y | y | y | y | y | y | y | y | y | n | y | y | y | y | y | y | y | y | y |
| Bacteria | sequenced | y | y | y | y | y | y | y | y | y | y | y | y | y | y | y | y | y | y | y | y | y | y | n | y | y | y | y | y | y | y | y | y |
| | post_normalization | y | y | y | y | y | y | y | y | y | y | y | y | y | y | y | y | y | y | y | y | y | y | n | y | y | y | y | y | y | y | y | y |
| Fungi | sequenced | y | y | y | y | y | y | y | y | y | y | y | y | y | y | y | y | y | y | y | y | y | y | n | y | y | y | y | y | y | y | y | y |
| | post_normalization | y | n | y | n | y | n | n | y | y | y | y | n | y | y | y | y | y | y | y | n | y | y | n | n | y | y | y | y | y | y | y | n |
| Archaea | sequenced | y | y | y | y | y | y | y | y | y | y | y | y | y | y | y | y | y | y | y | y | y | y | n | y | y | y | y | y | y | y | y | y |
| | post_normalization | y | n | y | n | y | y | y | y | y | y | y | y | y | n | y | y | y | y | y | y | y | y | n | y | y | y | y | y | y | y | n | y |

| | | P19 | | | | P20 | | | | P21 | | | | P22 | | | | P23 | | | | P24 | | | | P25 | | | | P26 | | | |
|---|---|---|---|---|---|---|---|---|---|---|---|---|---|---|---|---|---|---|---|---|---|---|---|---|---|---|---|---|---|---|---|---|---|
| | | O | U | V | S | O | U | V | S | O | U | V | S | O | U | V | S | O | U | V | S | O | U | V | S | O | U | V | S | O | U | V | S |
| Sample | collected | y | y | y | n | y | n | n | n | y | y | y | y | y | y | y | y | y | y | y | y | y | y | y | y | y | y | y | y | y | y | y | n |
| Bacteria | sequenced | y | y | y | n | y | n | n | n | y | y | y | y | y | y | n | n | y | y | y | y | y | y | y | n | y | y | y | y | y | y | y | n |
| | post_normalization | y | y | y | n | y | n | n | n | y | y | y | y | y | y | n | n | y | y | y | y | y | y | y | n | y | y | y | y | y | y | y | n |
| Fungi | sequenced | y | y | y | n | y | n | n | n | y | y | y | y | y | y | y | y | y | y | y | y | y | y | y | y | y | y | y | y | y | y | y | n |
| | post_normalization | y | y | y | n | y | n | n | n | y | y | y | n | y | y | y | n | y | y | y | y | y | y | n | n | y | y | y | n | y | y | y | n |
| Archaea | sequenced | y | y | y | n | y | n | n | n | y | y | y | y | y | y | y | y | y | y | y | y | y | y | n | y | y | y | y | y | y | y | y | n |
| | post_normalization | y | y | y | n | y | n | n | n | y | n | y | y | n | y | y | y | y | n | n | n | y | n | n | y | y | y | n | y | y | y | y | n |

| | | P27 | | | | P28 | | | | P29 | | | | P30 | | | | P31 | | | | P32 | | | |
|---|---|---|---|---|---|---|---|---|---|---|---|---|---|---|---|---|---|---|---|---|---|---|---|---|---|
| | | O | U | V | S | O | U | V | S | O | U | V | S | O | U | V | S | O | U | V | S | O | U | V | S |
| Sample | collected | y | y | y | y | y | y | y | y | y | y | y | y | y | y | y | y | y | y | y | y | y | y | y | y |
| Bacteria | sequenced | y | y | y | y | y | y | y | y | y | y | y | y | y | y | y | y | y | y | y | y | y | y | y | y |
| | post_normalization | y | y | y | y | y | y | y | y | y | y | y | y | y | y | y | y | y | y | y | y | y | y | y | y |
| Fungi | sequenced | y | y | y | y | y | y | y | y | y | y | y | y | y | y | y | y | y | y | y | y | y | y | y | y |
| | post_normalization | y | y | y | y | y | y | y | n | y | y | y | n | y | y | y | n | y | y | y | n | n | y | y | n |
| Archaea | sequenced | y | n | y | y | y | y | y | y | y | y | y | y | y | y | y | y | y | y | y | y | y | y | y | y |
| | post_normalization | y | n | y | y | y | y | y | n | y | y | y | y | y | y | n | y | y | y | y | y | y | y | y | y |

mpost

| | | P01 | | | | P02 | | | | P03 | | | | P04 | | | | P05 | | | | P06 | | | | P07 | | | | P08 | | | |
|---|---|---|---|---|---|---|---|---|---|---|---|---|---|---|---|---|---|---|---|---|---|---|---|---|---|---|---|---|---|---|---|---|---|
| | | O | U | V | S | O | U | V | S | O | U | V | S | O | U | V | S | O | U | V | S | O | U | V | S | O | U | V | S | O | U | V | S |
| Sample | collected | y | y | y | n | y | y | y | n | y | y | y | n | y | y | y | n | y | y | y | n | y | y | y | n | y | y | y | n | y | y | y | n |
| Bacteria | sequenced | y | y | y | n | y | y | y | n | y | y | y | n | y | y | y | n | y | y | y | n | y | y | y | n | y | y | y | n | y | y | y | n |
| | post_normalization | y | y | y | n | y | y | y | n | y | y | y | n | y | y | y | n | y | y | y | n | y | y | y | n | y | y | y | n | y | y | y | n |
| Fungi | sequenced | y | y | y | n | y | y | y | n | y | y | y | n | y | y | y | n | y | y | y | n | y | y | y | n | y | y | y | n | y | y | y | n |
| | post_normalization | y | n | y | n | y | y | y | n | y | y | y | n | y | y | y | n | y | y | y | n | y | y | y | n | y | n | n | n | y | y | y | n |
| Archaea | sequenced | y | y | y | n | y | y | y | n | y | y | y | n | y | y | y | n | y | y | y | n | y | y | y | n | y | y | y | n | y | y | y | n |
| | post_normalization | y | n | y | n | y | y | n | n | y | n | y | n | y | n | y | n | n | y | y | n | y | y | y | n | y | n | y | n | y | n | n | n |

| | | P09 | | | | P10 | | | | P11 | | | | P12 | | | | P14 | | | | P15 | | | | P16 | | | | P18 | | | |
|---|---|---|---|---|---|---|---|---|---|---|---|---|---|---|---|---|---|---|---|---|---|---|---|---|---|---|---|---|---|---|---|---|---|
| | | O | U | V | S | O | U | V | S | O | U | V | S | O | U | V | S | O | U | V | S | O | U | V | S | O | U | V | S | O | U | V | S |
| Sample | collected | y | y | y | n | y | y | y | n | y | y | y | n | y | y | y | n | y | y | y | n | y | y | y | n | y | y | y | n | y | y | y | n |
| Bacteria | sequenced | y | y | y | n | y | y | y | n | y | y | y | n | y | y | y | n | y | y | y | n | y | y | y | n | y | y | y | n | y | y | y | n |
| | post_normalization | y | y | y | n | y | y | y | n | y | y | y | n | y | y | y | n | y | y | y | n | y | y | y | n | y | y | y | n | y | y | y | n |
| Fungi | sequenced | y | y | y | n | y | y | y | n | y | y | y | n | y | y | y | n | y | y | y | n | y | y | y | n | y | y | y | n | y | y | y | n |
| | post_normalization | y | y | y | n | y | y | y | n | y | y | y | n | y | y | y | n | y | y | y | n | y | y | y | n | y | y | y | n | y | y | y | n |
| Archaea | sequenced | y | y | y | n | y | y | y | n | y | y | y | n | y | y | y | n | y | y | y | n | y | y | y | n | y | y | y | n | y | y | y | n |
| | post_normalization | y | y | y | n | y | y | y | n | y | n | y | n | y | y | y | n | y | y | y | n | y | y | y | n | y | y | y | n | y | y | y | n |

| | | P19 | | | | P20 | | | | P21 | | | | P22 | | | | P23 | | | | P24 | | | | P25 | | | | P26 | | | |
|---|---|---|---|---|---|---|---|---|---|---|---|---|---|---|---|---|---|---|---|---|---|---|---|---|---|---|---|---|---|---|---|---|---|
| | | O | U | V | S | O | U | V | S | O | U | V | S | O | U | V | S | O | U | V | S | O | U | V | S | O | U | V | S | O | U | V | S |
| Sample | collected | y | y | y | n | y | y | y | n | y | y | y | n | y | y | y | n | y | y | y | n | y | y | y | n | y | y | y | n | y | y | y | n |
| Bacteria | sequenced | y | y | y | n | y | y | y | n | y | y | y | n | y | y | y | n | y | y | y | n | y | y | y | n | y | y | y | n | y | y | y | n |
| | post_normalization | y | y | y | n | y | y | y | n | y | y | y | n | y | y | y | n | y | y | y | n | y | y | y | n | y | y | y | n | y | y | y | n |
| Fungi | sequenced | y | y | y | n | y | y | y | n | y | y | y | n | y | y | y | n | y | y | y | n | y | y | y | n | y | y | y | n | y | y | y | n |
| | post_normalization | y | y | y | n | y | y | y | n | y | y | y | n | y | y | y | n | y | y | y | n | y | y | y | n | n | y | y | n | y | y | y | n |
| Archaea | sequenced | y | y | y | n | y | y | y | n | y | y | y | n | y | y | y | n | y | y | y | n | y | y | y | n | y | y | y | n | y | y | y | n |
| | post_normalization | y | y | y | n | y | y | n | n | n | y | n | n | y | y | n | n | y | y | y | n | n | y | y | n | y | n | y | n | y | y | y | n |

| | | P27 | | | | P28 | | | | P29 | | | | P30 | | | | P31 | | | | P32 | | | |
|---|---|---|---|---|---|---|---|---|---|---|---|---|---|---|---|---|---|---|---|---|---|---|---|---|---|
| | | O | U | V | S | O | U | V | S | O | U | V | S | O | U | V | S | O | U | V | S | O | U | V | S |
| Sample | collected | y | y | y | n | y | y | y | n | y | y | y | n | y | y | y | n | y | y | y | n | y | y | y | n |
| Bacteria | sequenced | y | y | y | n | y | y | y | n | y | y | y | n | y | y | y | n | y | y | y | n | y | y | y | n |
| | post_normalization | y | y | y | n | y | y | y | n | y | y | y | n | y | y | y | n | y | y | y | n | y | y | y | n |
| Fungi | sequenced | y | y | y | n | y | y | y | n | y | y | y | n | y | y | y | n | y | y | y | n | y | y | y | n |
| | post_normalization | y | y | n | n | y | y | y | n | y | y | y | n | n | y | y | n | y | y | y | n | y | y | y | n |
| Archaea | sequenced | y | y | y | n | y | y | y | n | y | y | y | n | y | y | y | n | y | y | y | n | y | y | y | n |
| | post_normalization | y | y | y | n | n | y | y | n | y | n | y | n | y | y | y | n | y | y | y | n | y | y | n | n |
